# Interleukin-6 trans-signaling is a candidate mechanism to drive progression of human DCCs during periods of clinical latency

**DOI:** 10.1101/2020.05.28.121145

**Authors:** Melanie Werner-Klein, Ana Grujovic, Christoph Irlbeck, Milan Obradovic, Martin Hoffmann, Huiqin Koerkel-Qu, Xin Lu, Steffi Treitschke, Cäcilia Köstler, Catherine Botteron, Kathrin Weidele, Christian Werno, Bernhard Polzer, Stefan Kirsch, Miodrag Guzvic, Jens Warfsmann, Kamran Honarnejad, Zbigniew Czyz, Isabell Blochberger, Sandra Grunewald, Elisabeth Schneider, Gundula Haunschild, Nina Patwary, Severin Guetter, Sandra Huber, Stefan Buchholz, Petra Rümmele, Norbert Heine, Stefan Rose-John, Christoph A. Klein

## Abstract

Although thousands of breast cancer cells disseminate and home to bone marrow until primary surgery, usually less than a handful will succeed in establishing manifest metastases months to years later. To identify signals that support survival or outgrowth in patients, we profiled rare bone marrow-derived disseminated cancer cells (DCCs) long before manifestation of metastasis and identified IL6/PI3K-signaling as candidate pathway for DCC activation. Surprisingly, and similar to mammary epithelial cells, DCCs lacked membranous IL6 receptor expression and mechanistic dissection revealed IL6 trans-signaling to regulate a stem-like state of mammary epithelial cells via gp130. Responsiveness to IL6 trans-signals was found to be niche-dependent as bone marrow stromal and endosteal cells down-regulated gp130 in premalignant mammary epithelial cells as opposed to vascular niche cells. *PIK3CA* activation rendered cells independent from IL6 trans-signaling. Consistent with a bottleneck function of microenvironmental DCC control, we found *PIK3CA* mutations highly associated with late-stage metastatic cells while being extremely rare in early DCCs. Our data suggest that the initial steps of metastasis formation are often not cancer cell-autonomous, but also depend on microenvironmental signals.

## Introduction

In breast cancer, dissemination to distant sites precedes the clinical manifestation of metastasis by six to eight years in median, ranging from less than one year to more than 40 years ^1–3^. These clinical data derived from breast cancer growth kinetics and imaging studies are strongly supported by recent experimental evidence. Whereas dissemination from the primary site occurs preferentially in early tumor stages ^4–6^, specific mechanisms reduce cancer cell dissemination in anatomically and molecularly advanced stages ^5^. Furthermore, analysis of cancer growth kinetics suggests that not only cancer cells disseminate early, but also that micro-metastatic colony formation is initiated early. However, its manifestation may take considerable time ^1, 3, 7^. Such data are consistent with the observation that early DCCs often lack critical genetic and genomic alterations, which they need to acquire at the distant site in breast and other cancers. This process could explain the much longer clinical latency periods observed in humans as compared to mouse models ^5, 8, 9^ and be particularly relevant for cancers displaying late relapses such as hormone receptor positive breast cancer. However, early dissemination and prolonged clinical latency at distant sites raise questions about the identity and nature of signals conferring survival, genomic progression and outgrowth of DCCs over extended periods of time.

DCCs are extremely rare. They are detected at very low frequencies (1-2 DCCs per 10^6^ BM cells ^10, 11^) in bone marrow (BM) of about 30% of breast cancer patients with no evidence of manifest metastasis. Besides genomic studies, the assessment of the DCC phenotype has been limited to testing for selected antigens ^12^ and to anecdotal transcriptomic studies ^13, 14^. Moreover, spontaneous or transgenic mouse models, such as the PyMT- or Her2-driven models, do not generate bone metastases. Hence, there is no *in vivo* model available to study the spontaneous progression and genomic evolution from early BM infiltration to manifestation of bone metastasis. To unravel mechanisms operative during clinical latency periods, we interrogated transcript-derived pathway information from DCCs isolated from BM of breast cancer patients. Since early breast cancer DCCs often display close-to-normal genomes ^5, 15^, we used mammary epithelial cells isolated from reduction mammoplasties and available immortalized pre-malignant breast cancer cell lines as cellular models for functional testing of candidate mechanisms *in vitro*. We identified IL6 trans-signaling as pathway that (i) activates normal and pre-malignant cells, (ii) induces a proliferative stem/progenitor-like phenotype in mammary epithelial cells and (iii) whose activation in DCCs depends on regulatory niche cells in bone marrow. These data shed light onto the so far dark stage of early metastasis formation in patients. Moreover, it may inform about future ways to delay or prevent metachronous metastasis in patients whose breast cancer is diagnosed to be locally confined by standard clinical means.

## Results

### Early DCCs do not engraft in immunodeficient mice

For decades attempts to culture early DCCs, i.e. DCCs from non-metastasized M0-stage patients, have failed. Only anecdotal reports have been published that were not reproduced since then ^16–18^. We recently observed in melanoma that early DCCs failing to generate xenografts differ genomically from DCCs that successfully engrafted, indicating a causative role of genomic “maturation” for metastasis and xenograft formation. Specific alterations were identified that are closely linked to colonization success in mice and patients ^9^. We repeated these experiments for BM-derived breast and prostate cancer DCCs from early (M0) and advanced (M1) stages. Given the very low frequency of BM-DCCs (<10^-6^), we either injected CD45 depleted or EpCAM-enriched BM cells or generated and transplanted spheres as these have a higher engraftment-likelihood ^9^. In total, we tested 42 patient samples and different routes of application, including sub-cutaneous, orthotopic (site of origin), intra-femoral and intra-venous injection. We then assessed tumor formation at the cutaneous injection sites and metastatic spread to lungs or BM. BM-derived DCCs from M1-stage patients engrafted in two out of four cases. In contrast, early DCCs from 42 M0-stage patients did not establish xenografts (Fig. 1a, b), neither at the injection sites nor in the lungs (p = 0.006; Fisher’s exact test). We also explored the presence of minimal systemic cancer by testing for human cytokeratin (CK) or EpCAM-positive cells in murine BM. Interestingly, albeit DCCs of non-metastatic patients did not expand in mice, they survived in murine BM in 4 out of 42 cases. We detected human EpCAM^+^ or CK^+^ DCCs at a frequency of 1-5 DCCs/million BM cells 4-14 weeks after injection of CD45-depleted human BM cells (Fig. 1c). For one of these rare events we could not only prove human and epithelial, but also malignant origin by single cell copy number alteration (CNA) analysis (Fig. 1d).

**Figure 1:**
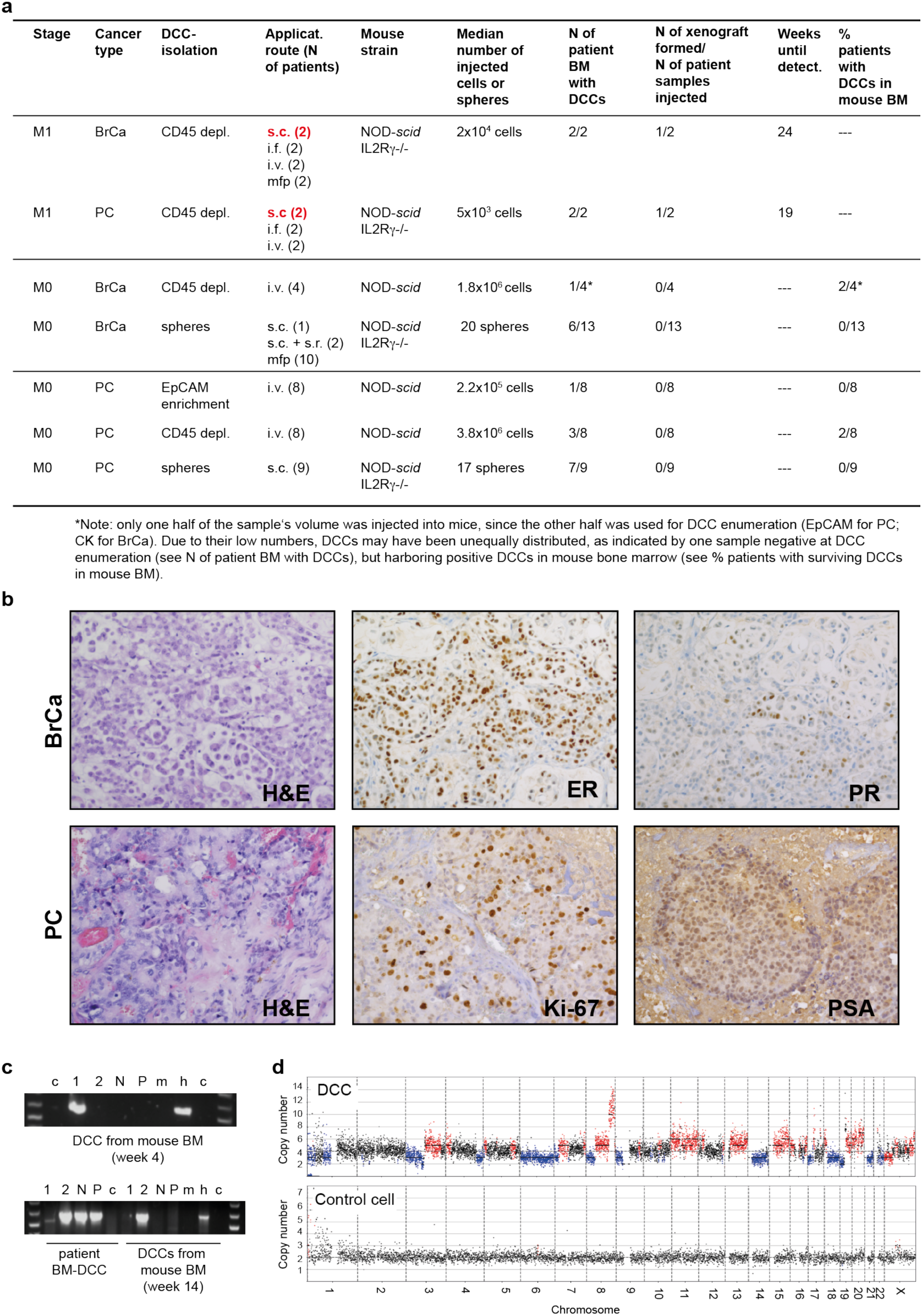
Xenotransplantation of DCCs. **a** Diagnostic bone marrow aspirates from breast (BrCa) or prostate (PC) cancer patients (M0- or M1- stage of disease) were either CD45-depleted, enriched for EpCAM or cultured under sphere conditions. Resulting spheres, CD45-depleted or EpCAM-enriched BM cells were injected intra-venously (i.v.), intra-femorally (i.f.), sub-cutaneously (s.c.), sub-renally (s.r.) or into the mammary fat pad (mfp) of NOD-*scid* or NOD-*scid*IL2Rg-/-mice. Mice with sub-cutaneous or mammary fat pad injections were palpated weekly. All other mice were observed until signs of illness or were sacrificed after 9 months. Injection routes that led to xenograft formation are highlighted in red. **b** Immunohistochemistry for estrogen-receptor (ER), progesterone-receptor (PR), prostate-specific antigen (PSA), Ki-67 or H&E staining of M1-DCC-derived xenografts is shown. **c** Human EpCAM- or cytokeratin 8/18/19-expressing DCCs were detected in the BM of 4/42 mice at 4 - 14 weeks after i.v. injection of CD45-depleted BM from non-metastasized patients. DCCs from two of the four mice were isolated and their human origin was verified by a PCR specific for human *KRT19*. Pure mouse or human DNA was used as control. 1, 2 =cytokeratin 8/18/19-positive DCCs; N=cytokeratin 8/18/19-negative BM-cell, P= pool of BM-cells of recipient mouse; m=mouse positive control; h=human positive control, c=non-template control. **d** Single cell CNA analysis of the EpCAM-expressing DCC isolated at 4 weeks after injection from NSG BM (panel c) and a human hematopoietic cell as control. Red or blue indicate gain or loss of chromosomal regions.

In summary and consistent with our findings in melanoma, early DCCs from patients without manifest metastasis failed to generate xenografts. Besides lower absolute cell numbers and fewer genetic alterations (see below), microenvironmental dependence of early DCCs could account for these results. We therefore decided to retrieve candidate interactions of early DCCs with the microenvironment via direct molecular analysis of early DCCs from breast cancer patients and implement these results into surrogate *in vitro* models.

### Pathway activation in mammary stem and progenitor cells

We hypothesized that stemness traits are necessary for the ability to survive and progress in a hostile environment and to initiate metastasis. Therefore, we tested for pathways activated in cells with progenitor or stem-like traits using our highly sensitive whole transcriptome amplification (WTA) method ^14, 19^. To identify these cells, we labeled freshly isolated primary human mammary epithelial cells (HMECs) from reduction mammoplasties of healthy patients with the membrane dye PKH26. Labelled cells were then cultured under non-adherent mammosphere conditions, which support the expansion of stem/early progenitor cells and formation of multicellular spheroids of clonal origin with self-renewing capacity ^20^. Cell divisions during mammosphere-formation diluted the dye until only few label-retaining cells (LRCs) were visible under the microscope (Fig. 2a). Isolating LRCs and non-LRCs (nLRCs) from disaggregated PKH26-labeled HMEC-spheres and plating them as single cell per well confirmed that the sphere-forming ability was solely confined to LRCs (Fig. 2b, Fisher’s exact test P=0.02), which is consistent with previous findings ^21^. For transcriptome analysis we isolated: (i) LRCs, (ii) nLRCs from disaggregated spheres and (iii) label-retaining cells that had not divided or formed spheres over time, but remained as single cells and were therefore termed quiescent single cells (QSCs). From each group we isolated single cells (LRC, n = 8; nLRC, n = 5; QSC, n = 10, Supplementary Table 1), performed WTA and microarray analysis as previously described ^19, 22^. Bioinformatics analysis indicated most variable gene expression between LRCs and QSCs/nLRCs (Fig. 2c, Supplementary Table 2) and we found twelve pathways significantly enriched in LRCs over QSCs/nLRCs (Fig. 2d, Supplementary Table 2, 3).

**Figure 2:**
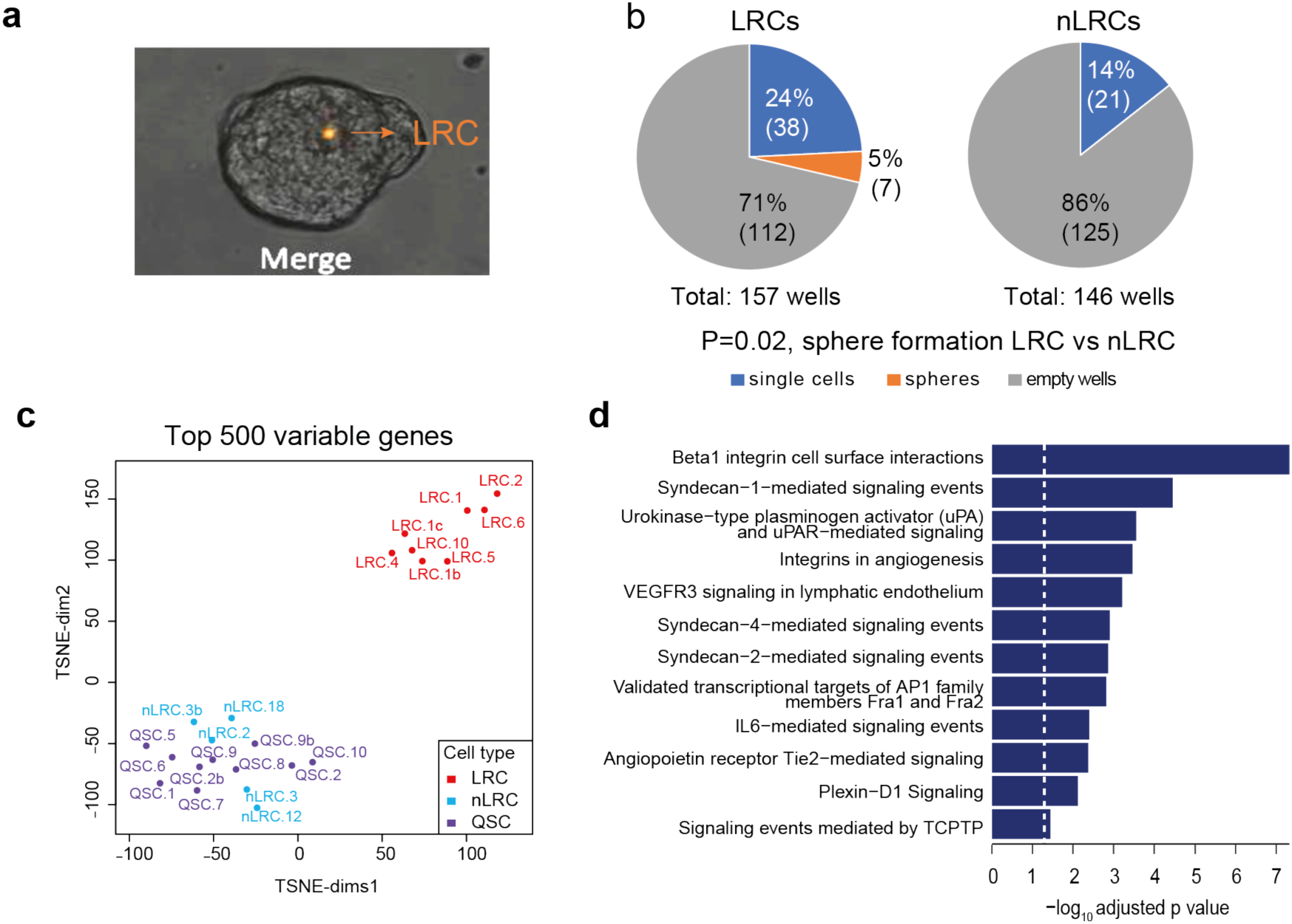
IL6 pathway is activated in mammary stem cells. **a** Representative picture of a mammosphere (day 7) generated from PKH26-labeled HMECs. **b** PKH26-labeled HMEC-mammospheres were disaggregated, sorted by flowcytometry into PKH26^+^ LRCs and PKH26^-^ nLRCs, plated as single cell per well and tested for sphere-formation. Shown is the percentage and in parentheses the respective absolute number of empty wells, wells with single cells and spheres after two weeks of mammosphere-culture (number of spheres vs. no spheres for LRCs vs. nLRCs, P=0.02, Fisher’s exact test). **c, d** LRCs (N=8), nLRCs (N=5) and QSCs (N=10) were subjected to single cell transcriptome microarray analysis. t-SNE plot of the top 500 most variable genes (panel c). Pathway analysis using the 216 genes differentially expressed between LRCs and the pooled nLRCs plus QSCs (panel d). See Supplementary Table 1 for patient/sample-ID allocation.

### Identification of EpCAM-positive DCCs in bone marrow

In order to test whether any of these pathways was enriched in DCCs isolated from BM of breast cancer patients, we aimed to isolate DCCs with confirmed malignant origin ^14^. We followed the reasoning that epithelial cell identity in bone marrow plus the presence of genetic alterations is sufficient to claim malignant epithelial origin or malignant potential of a cell. DCCs were detected by screening diagnostic BM aspirates from 246 M0-stage and 18 M1-stage patients for cells that stained positively for the epithelial marker EpCAM (Supplementary Fig. 1a). Forty percent of M0-stage and 72% of M1-stage patients harbored EpCAM^+^ cells. However, EpCAM is a surface marker that is not as specific for DCCs as the diagnostically used cytokeratins^14^ in bone marrow because it is expressed by cells from the B cell lineage (own unpublished data and ^23^). Therefore, we sought to differentiate between EpCAM-positive cells of breast cancer patients and non-cancer patients.

Copy number alterations (CNAs) are found in less than 5% of non-malignant cells with a median of 1.8% ^24–26^ and are diagnostically used to differentiate normal and malignant cells ^27^. We therefore performed combined genome and transcriptome analysis and isolated genomic DNA and mRNA from the same single cell ^5, 14^. Although this approach fails in 10-50% ^13, 14, 28^ and a CNA profile cannot be obtained for every cell, we found that 50% and 80% of successfully analyzed EpCAM-positive cells from M0 and M1 patients, respectively, harbored CNAs (Fig. 3a, Supplementary Fig. 1a). We selected these DCCs for single cell RNA-Seq analysis (M0: n=30 DCCs, 21 patients; M1: n=11 DCCs, 5 patients). To provide additional evidence that the aberrant EpCAM-positive cells are derived from a non-hematopoietic cell lineage, we compared them with autochthonous EpCAM-positive bone marrow cells. The latter were isolated from patients without known malignant disease undergoing hip replacement surgery. Of note, cancer patient-derived and non-cancer patient-derived EpCAM-positive cells could be clearly separated using the overall gene expression as well as epithelial and B-cell annotated genes – with the exception of one M0 cell, which was therefore excluded from further analysis (Fig. 3b). Moreover, many cells from M0-stage breast cancer patients strongly expressed the marker gene *KIT* (Fig. 3b), characteristic for mammary luminal progenitor cells ^29^ (see below). Together, copy number alterations and the epithelial, non-hematopoietic phenotype of cells isolated from patients with breast cancer provided compelling evidence that the selected cells were true DCCs.

**Figure 3:**
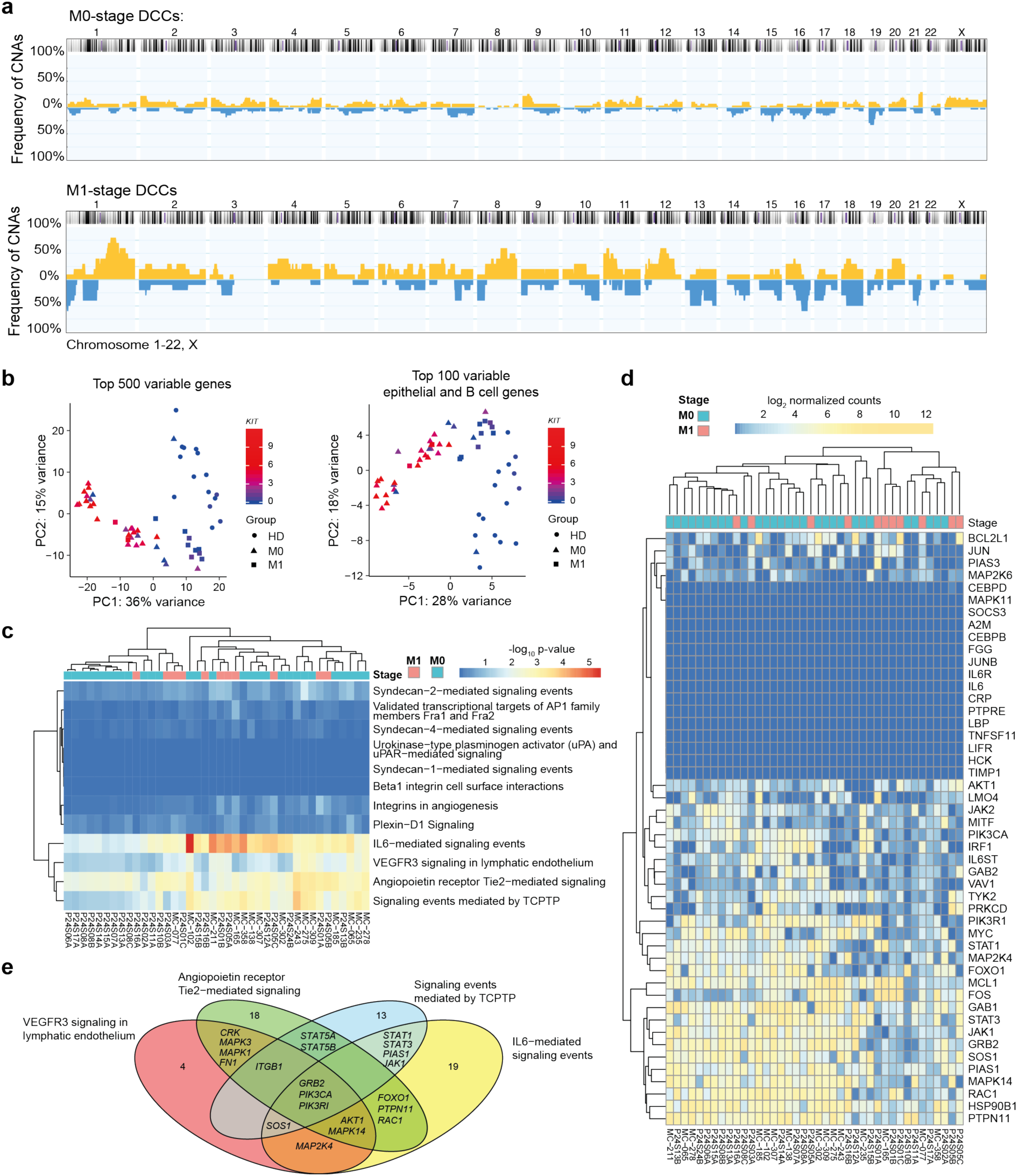
IL6 pathway is activated in BM-DCCs of breast cancer patients. **a** DCCs from BM of 21 non-metastasized (M0-stage, N=30 DCCs) and five metastasized (M1-stage; N=11 DCCs) breast cancer patients were analyzed for CNAs. The cumulative frequency of genomic aberrations is given in yellow and blue indicating genomic gains and losses, respectively. **b, c, d, e** M0- and M1-stage DCCs and EpCAM+ BM cells of seven healthy donors, i.e. patients without malignant disease (HD; N=15 cells) were analyzed by single cell RNA sequencing. Panel b: Principal component analysis of the top 500 most variable genes or top 100 most variable epithelial and B cell genes. Panel c: DCCs were tested for enrichment in pathways identified to be enriched in LRCs over QSCs/nLRCs (Fig. 2d). Panel d: The heatmap displays log2 normalized read counts of mRNA expression of IL6 signaling pathway genes as listed in the NCI-Nature PID expanded by the LIFR gene. Panel e: Venn diagram for the gene-members of the four pathways (panel d) that are expressed in at least half of bone marrow DCCs (except for the BMX (5/40) and CEBPD (19/40) genes, Supplementary Fig. 1c). Pathway-private genes are annotated by their number, shared genes are named explicitly (see also Supplementary Table 5). See Supplementary Table 1 for patient/sample-ID allocation.

### IL6 pathway activation in DCCs

We then tested whether any of the pathways enriched in mammary stem cells (LRCs, Fig. 2d) were also enriched in DCCs using pathway membership enrichment analysis. We found four out of the twelve pathways to be significantly enriched in DCCs (Fig. 3c, d, Supplementary Table 4) including the pathway “IL6-mediated signaling events”, the “TCPTP” pathway, the “VEGF-VEGFR3” and “Angiopoietin-Tie2 receptor” pathways. We decided to experimentally follow-up on the pathway “IL6-mediated signaling events” for several reasons: (i) IL6 signaling was previously found to be relevant for stemness maintenance, i.e. mammosphere-formation of ductal breast carcinoma and normal mammary gland ^30^; (ii) the TCPTP pathway, a negative regulator of IL6 signaling ^31^, was also enriched and (iii) assessment of individual genes expressed in these pathways (Supplementary Fig. 1c) revealed substantial overlap of the four pathways (Fig. 3e, Supplementary Table 5), indicating that related signaling modules had been triggered. We also tested the expression of the extracellular signal receptors and found that neither the receptor VEGFR3 nor Tie2 were expressed by DCCs. In contrast, while the mRNA of *IL6RA* (the IL6 binding receptor unit) was also absent in DCCs, the IL6 signal transducing unit gp130 was expressed (*IL6ST*, Fig. 3d) indicating amenability of DCCs to solely IL6 trans-signaling (see below) and thereby to microenvironmental control. Given the hints for a role of IL6 signaling for stemness maintenance and the restricted expression of IL6 signaling molecules, we decided to explore the activation of the IL6 pathway in normal and pre-malignant mammary cells in detail.

### IL6 trans-signaling activates sphere-forming ability

The IL6 pathway can be activated directly or *in trans*. Direct or classical IL-6 signaling involves IL6 binding to the heterodimeric receptor consisting of the ubiquitously expressed signal transducing receptor subunit gp130 and the membrane-bound IL6 receptor alpha chain CD126 ^32^. In contrast, trans-signaling does not involve the membranous IL6RA (mIL6RA), but binding of IL6 to the soluble IL6R alpha chain (sIL6RA) prior to binding to gp130 on the cell surface. sIL6RA can be generated by alternative splicing or limited proteolysis of the membrane-bound receptor and provided via autocrine and paracrine secretion. To explore the impact of the different modes of IL6 signaling on stemness or early progenitor traits, we used the pre-malignant human mammary epithelial cell lines MCF 10A and hTERT-HME1 and primary HMECs as models for early, genetically immature DCCs. Since metastasis founder cells are thought to display stem-like features ^33, 34^, cells were cultured under mammosphere conditions. Analysis of expression and secretion of IL6 signaling molecules by MCF 10A and primary HMECs using ELISA, flow cytometry and single cell PCR of LRCs and nLRCs (Supplementary Fig. 2a-d) indicated that (i) membranous IL6RA is expressed only in a fraction of LRCs and nLRCs; (ii) expression of IL6 signaling molecules (IL6RA, IL6, gp130) does not significantly differ between LRCs and nLRCs, (iii) co-expression of all signaling molecules in individual cells is extremely rare and (iv) the soluble form of IL6RA (sIL6RA) is generated by shedding of mIL6RA and not by splicing. Therefore, and in line with our results for breast cancer DCCs, IL6 trans-signaling via binding of IL6 to sIL6RA complexes to gp130 is much more likely involved in pathway activation than classical IL6 signaling. We therefore asked if stemness or early progenitor traits in mammary epithelial cells or DCCs are activated by a paracrine mode via classical signaling or trans-signaling. As a model for endogenous trans-signaling activation, we identified normal mammary cell-derived hTERT-HME1 cells with a knock-in of constitutively active EGFR (hTERT-HME1-EGFR^Δ746–750^) ^35^. This genetic change resulted in significantly increased amounts of IL6 trans-signaling components in the culture-supernatant (Supplementary Fig. 2e).

We then treated mammosphere cultures of MCF 10A, hTERT-HME1, hTERT-HME1-EGFR^Δ746–750^ and primary HMECs with activators or inhibitors of both pathways: (i) an anti-IL6 antibody to inhibit IL6 classical and trans-signaling, (ii) IL6 to activate classical and trans-signaling and (iii) Hyper-IL6 (HIL6) to selectively activate trans-signaling. HIL6 is a fusion protein consisting of sIL6RA, a linker chain, and IL6 and is used as a molecular model of the IL6/sIL6RA complex ^36, 37^. Adding IL6 or HIL6 to MCF 10A, hTERT-HME1 cells or HMEC cultures significantly increased sphere-formation (Fig. 4a-c, Student’s t-test P<0.01 or one-way ANOVA/Dunnett’s test P<0.0001). Interestingly, primary HMECs responded only to HIL6, but not IL6 (Fig. 4c) indicating that (i) the increase in sphere-number was due to IL6 trans-signaling, (ii) spheres originated from cells without mIL6RA expression and (iii) endogenous sIL6RA is a limiting factor (Supplementary Fig. 2c). Of note, hTERT-HME1-EGFR^Δ746–750^ could only marginally be stimulated by addition of HIL6, suggesting that it added little to the already available IL6/sIL6RA complexes (Fig. 4d).

**Figure 4:**
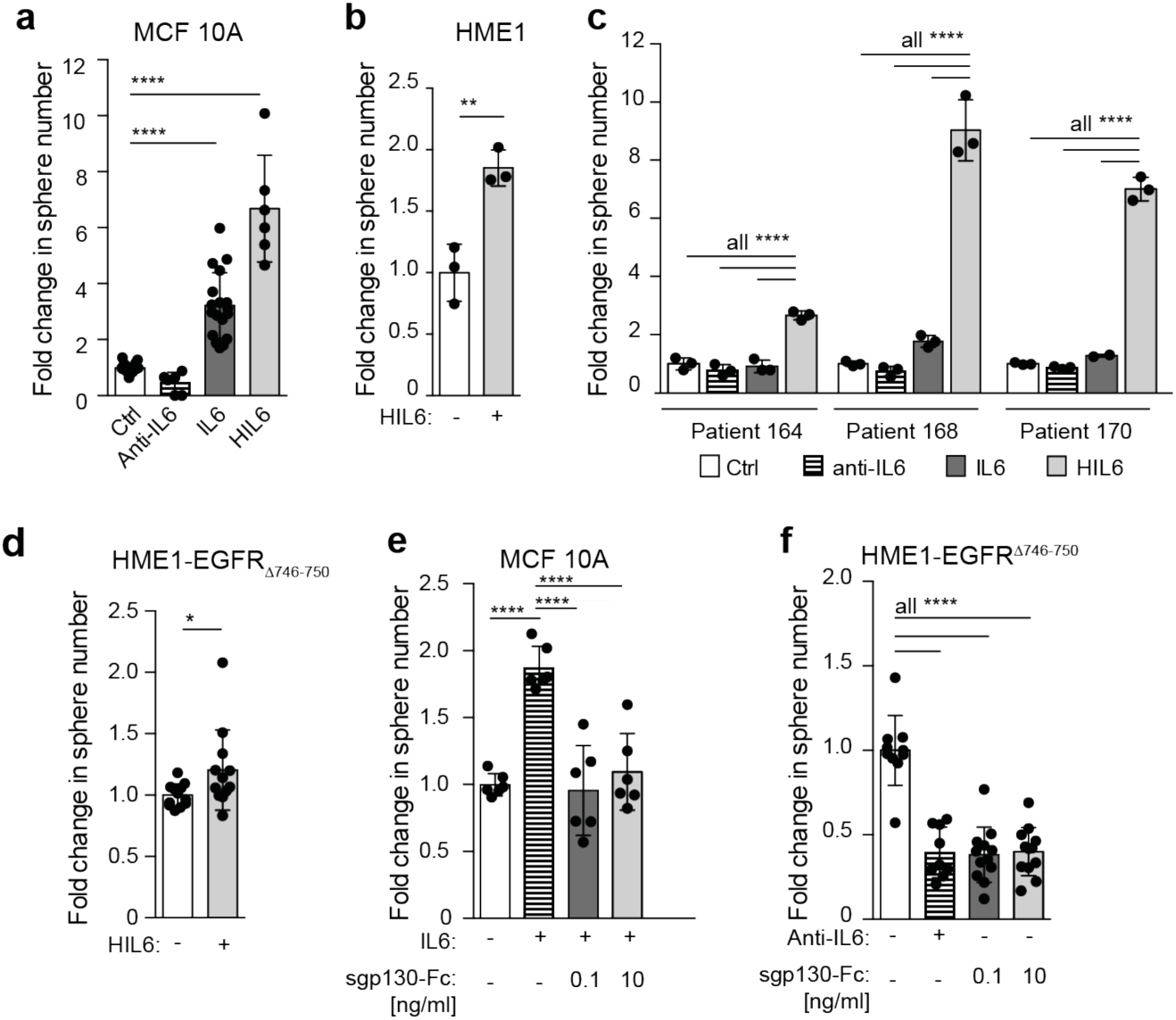
IL6 trans-signaling regulates the frequency of MCF 10A, hTERT-HME1 and primary HMECs with sphere-forming ability. **a** MCF 10A cells were cultured as spheres in the absence (N=18) or presence of IL6 (N=18), an IL6-blocking antibody (N=6) or Hyper-IL6 (N=6). **b** hTERT-HME1 were cultured as spheres in the absence (N=3) or presence (N=3) of Hyper-IL6. **c** HMECs were cultured without or with IL6, with an IL6 blocking antibody or Hyper-IL6. N=3 patients, each patient analyzed in triplicate. **d** hTERT-HME1-EGFR^Δ746–750^ cells were cultured as spheres in the absence or presence of HIL6 (each N=12). **e** MCF 10A cells were cultured as spheres without (N=6) or with IL6 (N=6) and IL6 plus sgp130-Fc at indicated concentrations (each N=6). **f** Sphere formation of hTERT-HME1-EGFR^Δ746-750^ in the absence (N=10) or presence of an anti-IL6 antibody (N=9) or with sgp130-Fc at indicated concentrations (each N=12). Cumulative data of three experiments. P values in panel a, c, f: one-way ANOVA with Dunnett’s multiple comparisons test (post hoc); panel b, d: two-sided Student’s t-test; panel e: one-way ANOVA with Tukey’s multiple comparisons test (post hoc); asterisks indicate significance between groups (*P<0.05, ** P<0.01, **** P<0.0001). All error bars correspond to standard deviation (Mean ± SD).

To dissect the impact of classical and trans-signaling on the observed increase in sphere-formation and hence the number of cells with stem-like activity, we specifically inhibited IL6 trans-signaling, but not classical signaling by adding the soluble form of gp130 (sgp130-Fc) to IL6 stimulated MCF 10A sphere-cultures ^38, 39^. At both concentrations of sgp130-Fc tested, IL6-induced sphere-formation was abolished (Fig. 4e; one-way ANOVA/Dunnett’s test P<0.0001) demonstrating that cells devoid of membranous IL6RA accounted for the increase in sphere-numbers by acquiring or activating stem-like functions in response to IL6 trans-signaling. Consistently, blocking of endogenous classical and trans-signaling in hTERT-HME1-EGFR^Δ746–750^ by anti-IL6, did not reduce sphere-formation to a greater extent than blocking IL6 trans-signaling only (Fig. 4f, one-way ANOVA/Dunnett’s test P<0.05 and <0.01).

### IL6 trans-signaling converts progenitor into stem-like cells

We noted that IL6- and HIL6-stimulated mammosphere-cultures showed an increase in the relative abundance of CD44^high^/CD24^low^ cells (Fig. 5a, b), a phenotype that has been ascribed to neoplastic and non-tumorigenic mammary cells enriched in tumor-initiating and sphere-forming cells, respectively ^33, 40^. Here, HIL6-stimulated cultures displayed the highest increase (Fig. 5a, b; one-way ANOVA/Dunnett’s test ctrl. vs. IL6 +/-sgp130-Fc, P<0.01; ctrl. vs. HIL6, P<0.0001). The increase in CD44^high^/CD24^low^ cells was not the result of increased proliferation of any CD24/CD44 subpopulation, but seemed to be caused by conversion of non-stem-like CD44^high^/CD24^high/int^ into CD44^high^/CD24^low^ stem-like cells (Supplementary Fig. 3a-c). To corroborate these findings, we compared IL6/HIL6-induced differential gene expression in MCF 10A cells to differences in gene expression between mammary stem cell enriched (MaSC), luminal progenitor (LumProg) and mature luminal (MatLum) cells as published by Lim et al. ^41^. The overlap between the respective differentially expressed genes was highly significant in almost all comparisons (Fig. 5c, Supplementary Fig. 3d, Supplementary Table 6) and the observed expression fold changes were consistent with the notion that IL6/HIL6 stimulation recruits progenitor populations from both, more differentiated as well as more stem-like populations, with the de-differentiation branch (MatLum → LumProg) being more consistent than differentiation (MaSC → LumProg). It should be noted that these *in vitro* generated data are fully consistent with the strong expression of the luminal progenitor marker *KIT* in DCCs (Fig. 3b) that we found to be activated via IL6 signaling.

**Figure 5:**
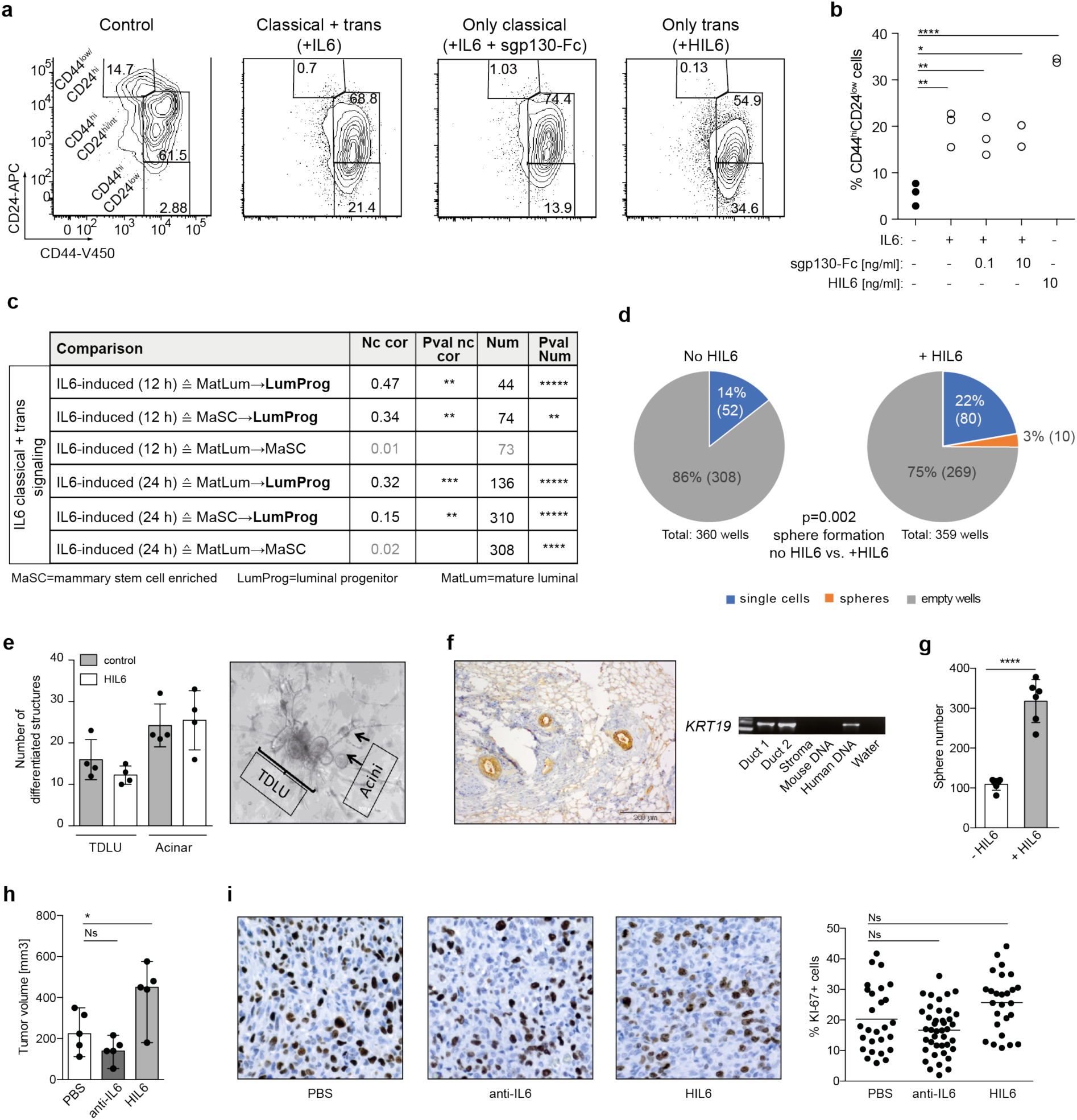
IL6 trans-signaling endows non-stem cells with stem-like abilities. **a**, **b** MCF 10A spheres were analyzed by flow cytometry for the expression of CD44 and CD24. The percentage of CD44^hi^CD24^low^ expressing cells was determined. Data represent cumulative results of three independently performed experiments, each performed in triplicate. **c** Fold change correlation analysis comparing IL6-induced gene expression in MCF 10A cells at 12 and 24 hrs, respectively, with gene expression signatures of luminal progenitor (LumProg), mature luminal (MatLum) and mammary stem cell enriched cells (MaSC) according to the study of Lim et al. ^32^. Nc cor: non-centered correlation between fold-changes, Num: number of common differentially expressed genes; **d** LRCs and nLRCs were sorted by flow cytometry from PKH26-labeled HMEC-spheres, nLRCs were plated as single cell per well (N=3 patients, single-cell deposit determined by manual microscopic evaluation) and cultured under mammosphere-conditions with or without HIL6. Sphere-formation and survival of single cells was determined after 14 days (P values are provided within panel d). Each patient-culture was set-up as duplicate in either freshly prepared or conditioned mammosphere-medium. As no significantly different outcome (Fisher’s exact test, P=0.6 and P=1 fresh vs. conditioned medium for cultures w/o HIL6 and with HIL6, respectively) was detected, results are presented as pooled analyses. **e** *In vitro* generation of acinar and tubular (TDLU) structures of primary HMECs cultured with or without HIL6 (each N=4). **f** Primary HMECs cultured with HIL6 and transplanted into NSG-mice. Staining for human cytokeratin 8/18/19 of the transplanted area eight weeks post-transplantation. PCR specific for human *KRT19* of two microdissected cytokeratin 8/18/19-positive ducts and one cytokeratin 8/18/18-negative stromal area. Pure mouse or human DNA was used as control. **g** MDA-MB-231 cells were cultured as spheres in the absence (N=6) or presence of HIL6 (N=6). **h** Tumor volume of 20,000 MDA-MB-231 cells pre-treated for 3 hours with PBS, an anti-IL6 antibody or HIL6 and transplanted into NSG-mice. All mice were analyzed at the same day after tumor cell inoculation. **i** TissueFAX cytometric quantification of tumors from panel h for the percentage of Ki-67-positive cells. N=33, 41 or 26 regions (0.25 mm^2^ each) for PBS, anti-IL6 or HIL6. P values in panel b, h, i: one-way ANOVA with Dunnett’s multiple comparisons test (post hoc); panel c: P values according to Student’s t-distribution for Nc cor and hypergeometric testing for Num; panel d: Fisher’s exact test; g: Student’s t test; asterisks indicate significance between groups in multiple comparisons (* P<0.05, ** P<0.01, **** P<0.0001). All error bars correspond to standard deviation (Mean ± SD).

To confirm these findings, we tested if *ex vivo* derived primary HMECs converted to stem-like cells by HIL6 activation. We isolated nLRCs from non-IL6-stimulated HMEC-mammospheres (Supplementary Fig. 3e) and re-plated them as single cell per well with or without HIL6. Whereas in the absence of HIL6 nLRCs were unable to form spheres, the proportion of sphere-forming cells induced from nLRCs in the presence of HIL6 was similar to that of non-HIL6 stimulated LRCs (3 % vs. 5 % sphere-formation see Fig. 5d and 2b). Moreover, replacing HIL6 by IL6 in the first or second week of a two-week mammosphere assay showed that continuous IL6 trans-signaling is needed to induce and maintain the number of cells with stem-like activity (Supplementary Fig. 3f). Finally, we confirmed that primary HIL6-treated HMEC spheres retain their ability to form acinar and tubular structures *in vitro* (Fig. 5e) and mammary ducts in immunodeficient NSG-mice (Fig. 5f). As MCF 10A cells do not form tumors in immunodeficient mice, we selected the luminal progenitor derived MDA-MB-231 cells to test whether the HIL6 induced increase in sphere-formation translates into higher malignancy *in vivo*. Like MCF 10A cells, MDA-MB-231 cells show increased sphere formation in response to HIL6 (Fig. 5g). Upon xenotransplantation of an equal number of MDA-MB-231 cells pre-treated with PBS, anti-IL6 or HIL6 for 3 hours, tumors in the HIL6-group were significantly larger than in the control group (Fig. 5h). This was not caused by increased proliferation or decreased apoptosis as the percentage of Ki-67-positive tumor cells did not differ significantly between the groups (Fig. 5i), and caspase-3 positive cells were not detected in any of the tumors.

### Bone marrow niche cells regulate responsiveness to IL6 trans-signaling

As gp130 expression is essential for IL6 signaling, we tested whether BM stromal cells modulate the ability of mammary epithelial cells to receive IL6 signals. We isolated primary human mesenchymal cells (MSCs) from diagnostic BM-aspirates of non-metastasized breast cancer patients or healthy volunteers and confirmed their ability to differentiate into adipocytes and osteoblasts *in vitro* (Fig. 6a, Supplementary Fig. 4a). We then co-cultured MCF 10A cells with (i) MSCs, (ii) *in vitro* differentiated osteoblasts (OBs) or (iii) human umbilical vein endothelial cells (E4ORF1-HUVECs ^42^) under non-sphere conditions. Interestingly, flow cytometric analysis revealed cell surface down-regulation of gp130 on MCF 10A cells co-cultured with MSCs and OBs, but not with HUVECs (Fig. 6b). Separation of MCF 10 A and MSCs by a transwell or using MSC-conditioned medium (CM) showed gp130 cell surface down-regulation to be independent from cell-cell contact (Supplementary Fig. 4b). Moreover, down-regulation was not immediate but observed between 6 and 14 hours after initiation of the co-culture with MSCs, OBs or MSC-conditioned medium from healthy donors or breast cancer patients (Fig. 6c, Supplementary Fig. 4c). This kinetic is consistent with the known independency of gp130-internalization from ligand binding ^43^ and points towards a transcriptional regulation of gp130 surface expression. To test this, we determined gp130 gene expression levels in single cells isolated from MCF 10A/MSCs and MCF-7/MSC co-cultures. Interestingly, both cell lines decreased their gp130 gene expression in response to MSCs (Fig. 6d), which is consistent with transcriptional regulation and demonstrates that early DCCs as well as more advanced cancer cells can respond to signals from neighboring cells.

**Figure 6:**
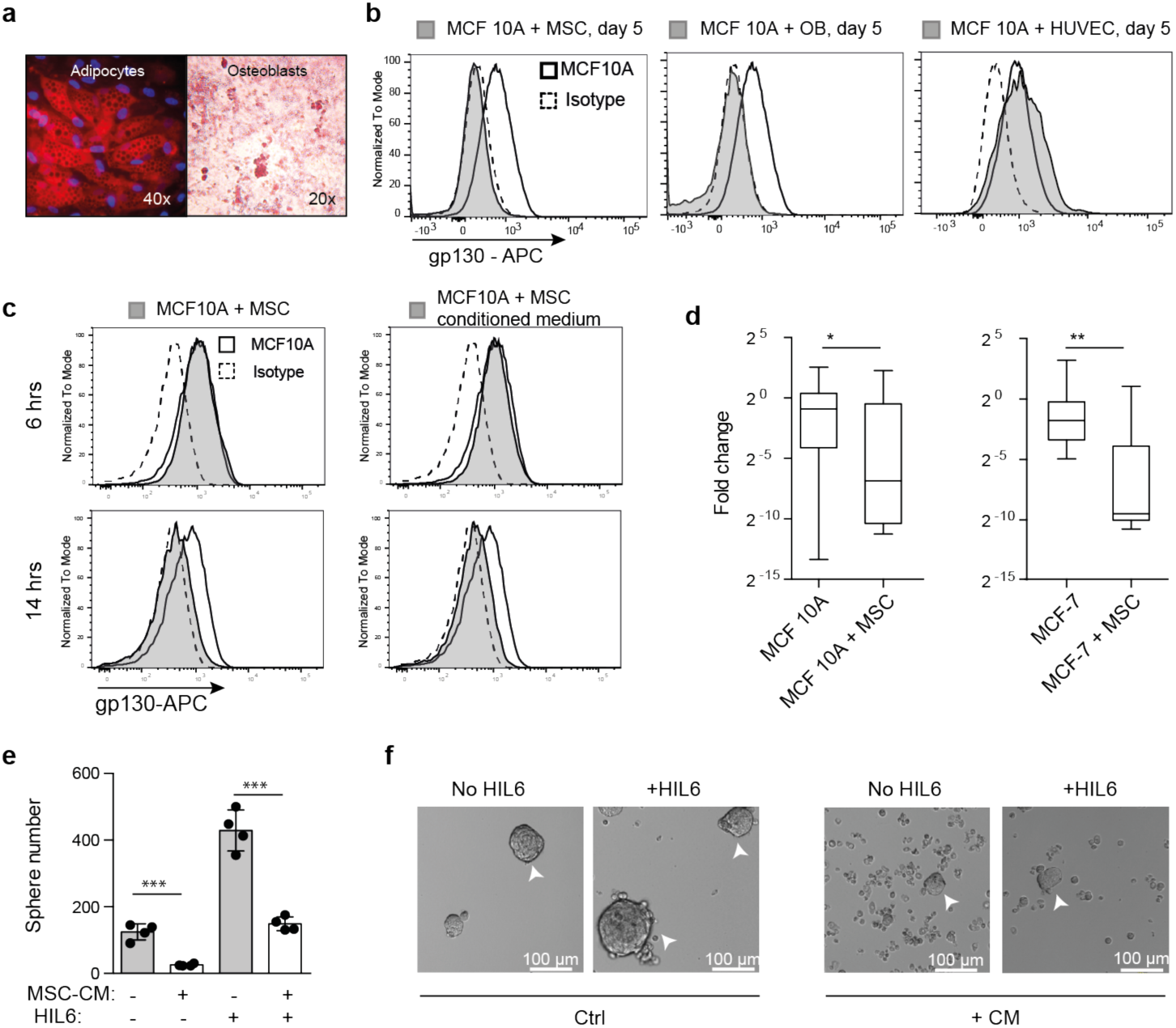
Surface expression of gp130 is down-regulated by soluble factors of bone marrow stromal cells. **a** Mesenchymal stem cells were tested for their ability to differentiate *in vitro* into adipocytes (Nil oil red O staining) and osteoblasts (Alizarin red S staining). **b** Surface gp130 expression of MCF 10A after five days of co-culture with primary mesenchymal stem cells (MSCs) from a breast cancer patient, primary osteoblasts (OBs) derived thereof or primary human umbilical vein endothelial cells (HUVECs). **c** Surface gp130 expression of MCF 10A after 6 and 14 hours of co-culture with MSCs or MSC-conditioned medium. Panel b, c: grey filled histograms indicate MCF 10A co-cultured with MSCs, OBs, HUVEC, MSC-conditioned medium or MSC separated by a transwell. Histograms with a thick black line indicate MCF 10A cells cultured alone and dashed histograms isotype control staining for gp130. **d** gp130 gene expression levels determined by single cell qPCR of MCF 10A cultured for 5 days with (N=25) or without (N=37) MSCs and MCF-7 cultured for 5 days with (N=20) or without (N=20) MSCs. Fold changes were calculated relative to MCF 10A or MCF-7 cells cultured without MSCs. **e**, **f** MCF 10A cells were left untreated or pre-treated for 14 hrs with MSC-conditioned medium, washed and then tested for their ability to form spheres in the presence of endogenously produced IL6/sIL6RA or exogenously added HIL6. Sphere-formation was assessed after seven days, N=4 for each group. P values according to two-sided Mann-Whitney test (panel d) or two-sided Student’s t-test (panel e). Asterisks indicate significance between groups **P<0.01,***P<0.001, ****P<0.0001). All error bars correspond to standard deviation (Mean ± SD).

To explore the functional impact of gp130 downregulation induced by MSCs, we tested MCF 10A cells pre-treated for 14 hrs with MSC-CM for their sphere-forming ability. Pre-treated MCF 10A showed a significant decrease in sphere-number and an increase in single, non-sphere forming cells in the presence of both, endogenously produced IL6/sIL6RA or exogenously added HIL6 (Fig. 6e, f, Student’s t-test, both P<0.001). The data indicate that the microenvironment in which early DCCs reside determines (i) their responsiveness to IL6 trans-signaling, with stromal and osteoblastic niches disabling IL6 trans-signaling in DCCs, and as a consequence (ii) the number of DCCs with stem-like phenotype and function, i.e. metastasis-initiating ability.

### Fully malignant DCCs escape IL6 trans-signaling dependence by oncogenic pathway activation

Cancer progression is driven by genetic and epigenetic evolution overriding microenvironmental control mechanisms. Although consensus about the nature of the metastatic niche is still lacking, MSC/OB-rich endosteal or vascular niches are believed to regulate the fate of DCCs ^42, 44^. Our experiments indicate that endosteal niches, although being rich in IL6 and sIL6RA molecules ^45–48^, render DCCs unresponsive to IL6 trans-signals. However, gp130^+^ DCCs in vascular niches would respond to IL6/sIL6RA complexes. Therefore, in both niches pathway activation by mutation would provide a selection advantage for DCCs that otherwise might depend on microenvironmental signals. We consequently sought for corroborating evidence that genetically variant DCCs may evade the need for IL6 trans-signaling and become selected. We considered the *PIK3CA* pathway a strong candidate for such a selected oncogenically activated pathway ^49^ as (i) IL6 signaling activates not only the JAK/STAT pathway, but also the PI3K/AKT pathway ^50^, (ii) early DCCs expressed *PIK3CA* as core element of the four identified stemness-associated pathways (Fig. 3e), (iii) activating *PI3KCA* mutations in exons 9 and 20 are among the most frequent mutations occurring in human breast cancer and (iv) constitutive activation of the *PIK3CA* pathway has been shown to evoke cell de-differentiation of mammary gland cells into a multipotent stem-like state ^51^. To test *PIK3CA*-signaling, we analyzed sphere-formation in response to HIL6 in pre-malignant MCF 10A and malignant breast cancer cell lines with or without a *PIK3CA* activating mutation. Interestingly, HIL6 increased sphere-formation only in cells with wildtype *PIK3CA* (Fig. 7a), but not in cells with an activating *PIK3CA^E545K/+^* mutation. Using isogenic MCF 10A cells with (knock-in of *PIK3CA^E545K/+^*) and without *PIK3CA* mutation, we noted that HIL6 activated the pSTAT3 pathway in mutant and wildtype cells (Fig. 7b, Supplementary Fig. 5). In contrast, IL6 trans-signaling induced a massive activation of the pERK and pAKT pathways only in wildtype cells as these pathways were already activated in PI3K-mutant cells (Fig. 7b). Consistently, untreated PI3K-mutant cells formed significantly more spheres than wildtype *PIK3CA* cells (Fig. 7c, Student’s t-test, P<0.0001). MSC-induced gp130 downregulation was unaffected by the *PIK3CA* mutational status (Supplementary Fig. 4d). In summary, these data show that *PIK3CA* activation overrides regulation of stemness-traits by IL6 trans signaling and renders cancers cells more independent from microenvironment control.

**Figure 7:**
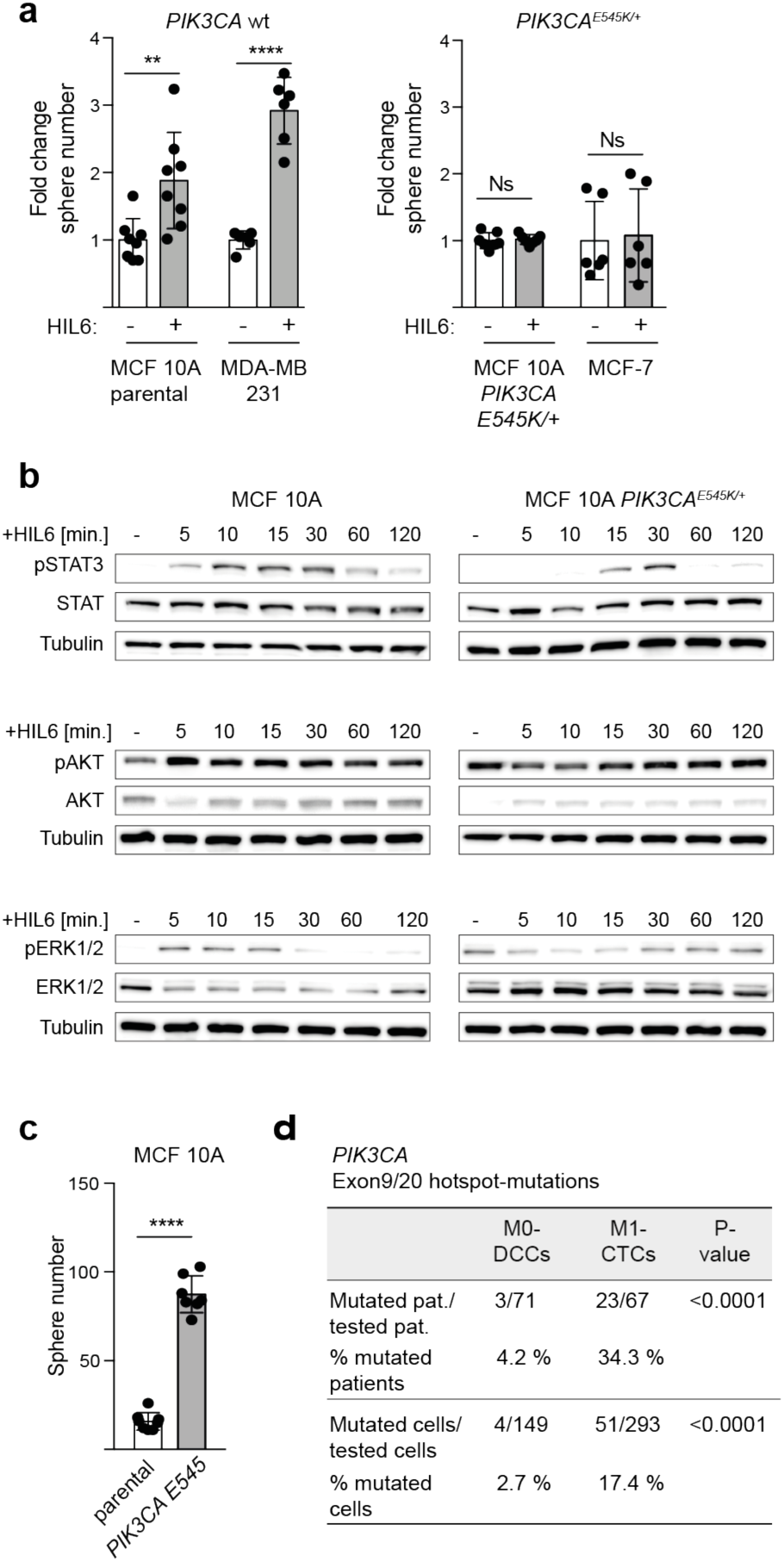
Activation of the *PIK3CA* pathway renders mammary epithelial cells independent of IL6 signaling. **a** Fold change in sphere numbers of pre-malignant (MCF 10A) and tumorigenic cell lines (MCF-7, MDA-MB-231) without (MCF 10A parental, N=8; MDA-MB-231, N=6) or with mutational activation of *PIK3CA* (MCF 10A *PIK3CA E545K/+,* N=7; MCF-7, N=6) cultured in the presence or absence of HIL6. Note that MCF 10A *PIK3CA E545K/+* cells are isogenic to MCF 10A parental. **b** Western blot analyses showing phosphorylation of STAT3^Tyr705^, AKT^Ser475^ and ERK1/2^Thr202/Tyr204^ in MCF 10A or MCF 10A *PIK3CA E545K/+* cells cultured without or with HIL6 for indicated time points. **c** Sphere numbers of the isogenic cells MCF 10A parental (N=8) and MCF 10A *PIK3CA E545K/+* (N=7) cultured in the absence of HIL6. **d** Cytokeratin 8/18/19^+^ DCCs from BM of non-metastasized (M0-stage) HR-positive breast cancer patients and CD45^-^/EpCAM^+^/cytokeratin 8/18/19^+^ CTCs isolated from peripheral blood of metastasized (M1-stage) HR-positive breast cancer patients were sequenced for hotspot-mutations in *PIK3CA* (Exon 9: E545K, E542K; Exon 20: H1047R, H1047L, M1043I). P values in panel b, c: two-sided Student’s test; d: Fisher’s exact test; asterisks indicate significance between groups (** P<0.001 and **** P<0.0001). All error bars correspond to standard deviation (Mean ± SD).

Based on these findings, we tested for direct evidence of genetic selection in the *PIK3CA* pathway during cancer progression by analyzing the *PIK3CA* gene for mutations in exon 9 and 20 in BM-derived DCCs from non-metastasized (M0-stage) and in circulating tumor cells (CTCs) from metastasized (M1-stage) breast cancer patients mostly displaying bone metastases. Both groups of cytokeratin-positive cancer cells had previously been shown to display CNAs ^5, 52^. As hormone-receptor positive breast cancer is the most frequent breast cancer type, we focused on this disease type to obtain meaningful sample numbers. Consistent with previous data, *PIK3CA* hotspot mutations were found in DCCs/CTCs of patients with hormone receptor (HR)-positive tumors (Fig. 7d). Strikingly, only 4.2% of M0-stage DCCs harbored *PIK3CA* mutations, whereas manifest metastasis-derived cells from HR-positive tumors displayed them in 34.3% of cases (Fisher’s exact test, P<0.0001, Fig. 7d). These data are fully consistent with a scenario in which early DCCs depend on IL6 trans-signaling and become increasingly independent thereof during cancer evolution.

## Discussion

In this study we provide evidence for a role of the niche microenvironment to enable and drive the earliest stages of human metastasis formation. We identified IL6 trans-signaling as an activator of stem-like and progenitor traits underlying epithelial colony formation, a mechanism that is characterized by dependence on both IL6 and sIL6RA, in contrast to IL6 alone, when a cell is equipped with mIL6RA. Our finding questions the concept of fully-malignant and autonomous cancer cells as founders of metastasis. However, during subsequent malignant evolution DCCs may evade microenvironmental control by acquiring IL6-independence. Our data indicate that this could occur via mutational activation of the PI3K pathway.

Several observations characterize early metastatic bone marrow colonization. First, breast cancer dissemination in humans often starts from lesions often measuring less than 1-4 mm in diameter ^2, 5^. Second, initially DCCs often do not display the typical karyotypic changes of breast cancer tumors or metastases, however are detected long before manifestation of clinical metastasis ^5, 8, 15^. Third, metastasis formation in BM usually takes years to decades ^53^ indicating that evolutionary mechanisms and slow growth kinetics dictate the pace. This framework precludes the use of transgenic mouse models that are too short-lived to mimic the process in patients and rarely form bone metastases. The same applies to *in vitro* and *in vivo* studies involving commonly used and genomically highly aberrant breast cancer cell lines derived from manifest metastases or primary tumors. All of these do not represent the biology under investigation.

We therefore aimed at expanding early DCCs in immune-deficient mice, but failed. Reasons for this can be manifold: DCCs are extremely rare and we estimate that often less than 10 cells/mouse were injected. Also, species barriers may preclude engraftment of cancer cells that may critically depend on certain microenvironmental signals, among them the species specific IL6 signaling ^54^. Our failure is also consistent with data from melanoma where we had identified that engraftment of DCCs requires activation of specific oncogenic pathways ^55^ - which in case of *PIK3CA* mutations early (M0-stage) breast cancer DCCs commonly lack. Consistently, M1-stage DCCs formed xenografts in two out of four cases. We therefore based our approach on two starting points: First, we analyzed transcriptomic data of early human breast cancer DCCs to identify candidate pathways and second, we used genomically close-to-normal and normal mammary epithelial cells as models to functionally interrogate the transcriptome data.

We focused our analysis on EpCAM-positive cells from bone marrow. Currently, EpCAM is the best marker to identify and isolate viable epithelial cells from bone marrow. However, it is neither fully specific ^14^ as cells of the B cell lineage may express EpCAM ^23^ nor does it identify all breast cancer initiating cells ^56^. However, EpCAM-positive mammary progenitor cells give raise to chromosomally instable cancers, such as triple negative basal like / post-EMT and hormone receptor positive cancers ^57^. Since we used chromosomal instability as inclusion criterion to identify DCCs our findings do not apply to claudin-low cancers which are derived from EpCAM-negative, chromosomally stable cells. Interestingly, early M0-stage DCCs isolated from bone marrow highly expressed *KIT*, which is characteristic for luminal progenitor cells. Luminal progenitor cells are the last common precursor of triple negative, basal-like and hormone receptor positive cancers ^58^. Since hormone-receptor positive breast cancers can experimentally be “normalized” by functionally impairing their luminal progenitor cells ^59^, cells with this phenotype are likely to comprise metastasis founder cells of EpCAM-positive triple negative and hormone-receptor positive cancers.

This reasoning is fully consistent with our observed effects of IL6 on normal mammary stem and progenitor cells. We identified an activated IL6 pathway in DCCs and our *in vitro* models revealed that IL6 trans-, but not classical signaling, induces de-differentiation of mammary epithelial cells and endows them with stemness and progenitor traits. IL6 trans-signaling makes cells dependent on the microenvironment, however, BM represents an IL6 rich environment ^47^. There, IL6/sIL6RA is important for regulation of hematopoiesis ^60^ and for generation of hematopoiesis-supporting BM stromal cells ^61^ using autocrine/paracrine feedback loops ^48^. In the context of breast cancer, it is noteworthy that serum levels of IL6 and sIL6R and their local production in BM by osteoblasts depend on sex-steroids, change with menstrual cycle, are negatively regulated by estrogen and hormone replacement therapy and increase up to tenfold during menopause ^48, 62, 63^. Therefore, systemic microenvironmental changes may provide a mechanism by which DCCs become activated in post-menopausal breast cancer patients. Consistently, bone-only metastasis is significantly associated with higher age at primary diagnosis in HR-positive breast cancer patients ^64^, known for very late relapses.

We show that IL6 trans-signaling equips normal and transformed mammary cells with stemness and progenitor traits likely to be crucial for DCCs. Although about 6,000 – 27,000 DCCs may lodge in the BM compartment ^65^, only few of them will progress to manifest metastasis. Our data are in line with this observation as co-culture with BM niche cells like osteoblasts or stromal MSCs resulted in loss of gp130 expression rendering only DCCs in vascular niches possibly responsive to IL6 trans-signaling. Moreover, only about 3-5% of mammary cells acquired mammosphere-formation ability upon HIL6 stimulation. Genetic analysis of DCCs revealed that DCCs from HR^+^ breast cancer progressing to manifest metastasis often acquired *PIK3CA* mutations, possibly because the activated PI3K/AKT pathway rendered mammary epithelial cells independent from IL6 trans-signaling.

We are aware of data indicating that some metastases may be derived from late-disseminating cells ^66, 67^. One study showed that in 3 out of 6 cases tested cytokeratin-positive cells displayed alteration profiles very similar to their matched primary tumors ^66^. Future studies will need to carefully address the role of tumor (sub)type, disease stage and duration, growth kinetics and a possible interdependence of early and late DCCs for metastasis formation. However, the findings presented here demonstrate that early DCCs are able to interpret signals from the bone marrow environment that activate them and might drive their progression. Thereby, our study may lay the groundwork for novel adjuvant therapies. Early DCCs striving to form metastasis during yearlong evolutionary processes may be sensitive to different drugs than fully malignant cancer cells. Their Achilles’ heel may consist of microenvironmental signals supporting their survival and genomic progression. Since there is hope that oncogene- or tumor-suppressor-gene-associated drug-resistance mechanisms are not yet operative, depriving early DCCs of microenvironmental support pathways could render them vulnerable to non-genotoxic drugs.

## Acknowledgements

We thank Irene Nebeja, Irina Hartmann, Michaela Becker and Silvia Materna-Reichelt for excellent technical assistance. We are grateful to Alberto Bardelli for hTERT-HME1 cell lines and Cyrus Ghajar for E4ORF1 HUVECs. This work was supported by grants to CAK from the Deutsche Krebshilfe (TransLUMINAL-B 111536), the Dr Josef Steiner foundation, the ERC (grant 322602), the Bavarian ministry of economic affairs, energy and technology (AZ 20-3410.1-1-1), the Deutsche Forschungsgemeinschaft (DFG; KL 1233/10-2) and the Bavarian Research Foundation (Bayerische Forschungsstiftung, DOK-165-13). The work of MW-K has been funded by the DFG (WE 4632/1-1, WE 4632-4/1 and WE 4632/5-1within the FOR2127). TissueFAX was supported by the DFG (INST 89/341-1 FUGG). The work of SR-J has been funded by grants from DFG in the SFB841 (project C1), the SFB 877 (projects A1 and A14), by the Deutsche Krebshilfe and by the German-Israeli Foundation for Scientific Research and Development.

## Authors’ Contributions

### Conception and design

C. A. Klein and M. Werner-Klein

### Development of methodology

M. Werner-Klein, A. Grujovic, M. Obradovic, M. Hoffmann, S. Kirsch, S. Treitschke, J. Warfsmann, K.Honarnejad, S. Rose-John

### Acquisition of data

M. Werner-Klein, A. Grujovic, C. Irlbeck, M. Obradovic, S. Treitschke, C. Köstler, C. Botteron, K. Weidele, C. Werno, S. Kirsch, M. Guzvic, K. Honarnejad, I. Blochberger, S. Grunewald, E. Schneider, G. Haunschild, N. Patwary, S. Guetter, S. Huber, S. Bucholz, P. Rümmele, N. Heine

### Analysis and interpretation of data

M. Werner-Klein, A. Grujovic, M. Hoffmann, X. Lu, H. Koerkel-Qu, B. Polzer, J. Warfsmann, K.Honarnejad, Z. Czyz, C. A. Klein

### Writing, of the manuscript

M. Werner-Klein, M. Hoffmann and C. A. Klein

### Review and/or revision of the manuscript

all authors

### Competing interest

The authors declare no potential conflicts of interest.

## Methods

### Patient material

Human non-cancerous mammary tissue was obtained from female patients undergoing reduction mammoplasty surgeries at the University Center of Plastic-, Aesthetic, Hand- and Reconstructive Surgery, University of Regensburg, Germany after informed, written consent of patients was obtained (ethics vote number 07/043, ethics committee of the University Regensburg). After verification of the non-cancerous origin of the tissue by a pathologist, mammary glands were dissociated and primary human mammary epithelial cells (HMECs) isolated.

Human disseminated cancer cells were obtained from BM-aspirates of breast or prostate cancer patients without and with distant metastases. EpCAM+ cells were obtained from bone marrow of patients without known malignant disease undergoing hip replacement surgery. Human mesenchymal stem cells were obtained from BM-aspirates of breast cancer patients or healthy donors. Written informed consent of cancer and control patients was obtained and the ethics committee of the University of Regensburg (ethics vote number 07/79) approved BM-sampling and analysis of isolated cells.

Enrichment and detection of CTCs was performed within the SUCCESS (EUDRA-CT number 2005-000490-21) and DETECT (EUDRA-CT number 2010-024238-46) ^68^ studies using the CellSearch® system ^69^. Written informed consent for CTC analysis and characterization was obtained for all patients included. All experiments conformed to the principles set out in the WMA Declaration of Helsinki and were approved by the ethical committees responsible for the corresponding studies (Universities of Munich, Dusseldorf, Tuebingen, and Ulm). Isolation and molecular analysis of CTCs was approved by the ethics committee of Regensburg (ethics vote number 07/079).

### Mice

NOD.Cg-Prkdc^scid^ IL2rg^tmWjl^/Sz (also termed NSG) or NOD.Cg-Prkdc^scid^ mice were purchased from the Jackson Laboratory, USA and maintained under specific-pathogen free conditions, with acidified water and food *ad libitum* in the research animal facilities of the University of Regensburg, Germany. All approved experimental animal procedures were conducted according to German federal and state government of Upper Palatinate, 54-2531.1-10/07, 54-2532.1-34/11; 54-2532.1-17/11, 54-2532.1-1/12, 54-2532.4-7/12).

### Cell lines

MCF-7 and MDA-MB-231 breast cancer cell lines were obtained from the German Collection of Microorganisms and Cell Cultures (DSMZ) and Cell Lines Service (CLS), respectively. MCF 10A (CRL-10317), a non-tumorigenic mammary epithelial cell line, was obtained from the American Type Culture Collection (ATCC^70^). The isogenic cell line MCF 10A PIK3CA^E545K/+^ (HD 101-002) together with its parental cell line MCF 10A-parental (HD PAR-003) were purchased from Horizon Discovery, United Kingdom. MCF 10A-GFP cells were generated by transducing MCF 10A cells with pRRL.sin.cPPT.hCMV-GFP.WPRE (generously provided by Luigi Naldini, Italy). hTERT-HME1-derived cell lines, E4ORF1-transduced primary human umbilical vein endothelial cells (HUVECs) and murine embryonic fibroblasts C3H10T1/2 were generously provided by Alberto Bardelli, (University of Turin, Italy), Cyrus Ghajar (Fred Hutchinson Cancer Research Center, USA) and Max Wicha (University of Michigan, USA), respectively. The identity of all cell lines was confirmed by DNA fingerprinting analysis utilizing the GenePrint 10 System (Promega).

All MCF 10A-derived cell lines were cultured in Ham’s Dulbecco’s modified Eagle’s/F12 (DMEM/F12) medium (Pan-Biotech, Germany) supplemented with 5% horse serum (Sigma-Aldrich, Germany), 2mM L-glutamine (Pan-Biotech, Germany), 1% penicillin/streptomycin (Pan-Biotech, Germany), 20 ng/ml EGF (Sigma-Aldrich, Germany), 0.5 μg/ml hydrocortisone (Sigma-Aldrich, Germany), 10 μg/ml insulin (Sigma-Aldrich, Germany) and 0.1 µg/ml cholera toxin (Sigma-Aldrich, Germany). All hTERT-HME1-derived cells lines were maintained in DMEM/F12 medium (Pan-Biotech, Germany) supplemented with 10% FCS (Sigma-Aldrich, Germany), 2mM L-glutamine (Pan-Biotech, Germany), 1% penicillin/streptomycin (Pan-Biotech, Germany), 20 ng/ml EGF (Sigma-Aldrich, Germany), 0.5μg/ml hydrocortisone (Sigma-Aldrich, Germany) and 10 μg/ml insulin (Sigma-Aldrich, Germany). MDA-MB-231 cells were cultured in DMEM medium (Pan-Biotech, Germany) supplemented with 10% FCS (Sigma-Aldrich, Germany), 2 mM L-glutamine (Pan-Biotech, Germany) and 1% penicillin/streptomycin (Pan-Biotech, Germany). MCF-7 cells were propagated in RPMI 1640 medium (Pan-Biotech, Germany) supplemented with 10% FCS, 2 mM L-glutamine and 1% penicillin/streptomycin. Murine embryonic fibroblasts C3H10T1/2 were grown in DMEM (Pan-Biotech, Germany) medium supplemented with 5% fetal calf serum (Pan-Biotech, Germany), 2mM L-glutamine (Pan-Biotech, Germany), 1% penicillin/streptomycin (Pan-Biotech, Germany). E4ORF1-transduced primary human umbilical vein endothelial cells (HUVECs) were cultured using the EGM-2 Bullet Kit (Lonza, Germany). All cell lines were kept at 37°C and 5% CO2 in a fully humidified incubator and negatively tested for mycoplasma by PCR.

### Isolation of disseminated cancer cells from bone marrow

Mononuclear cells from bone marrow of non-metastasized breast cancer patients were plated on adhesive slides (Thermo Fisher) at a density of 0,5-1×10^6^ cells/slide. Slides were stored at −20°C. From each patient, 1–2 10^6^ bone-marrow cells were stained with the monoclonal antibody A45-B/B3 (AS Diagnostik, Germany) against cytokeratin 8/18/19 and developed with the anti-mouse AB-Polymer (Zytomed Systems, Germany). Unspecific binding was blocked using PBS/10% AB-serum (Bio-Rad, Germany). Alkaline phosphatase was developed with 5-bromo-4-chloro-3-indolyl phosphate and Nitroblue tetrazolium (BCIP/NBT; BioRad, Germany) as substrate. Slides were covered with phosphate-buffered saline under a cover glass and assessed by bright-field microscopy. An identical number of cells served as a control for staining with mouse IgG1 Kappa (MOPC-21) without known binding specificity. After removal of the cover glass, positive cells were isolated from the slide with a micromanipulator (Eppendorf PatchMan NP2) and subjected to whole genome amplification for subsequent *PIK3CA* mutation analysis.

For isolation of both, RNA and DNA from the same disseminated cancer cells, mononuclear cells from BM of non-metastasized breast cancer patients were subjected to immunofluorescent staining for EpCAM (Ber-EP4-FITC, Agilent or HEA-125-PE, Miltenyi Biotec, Germany). Positive cells were isolated with a micromanipulator (Eppendorf PatchMan NP2, Eppendorf, Germany) and cells were subjected to whole transcriptome amplification to isolate RNA for subsequent PCR-analyses, transcriptome microarrays or RNA-seq and whole genome amplification for isolation of genomic DNA for subsequent analysis of copy number alterations.

### Isolation of circulating tumor cells

Up to three 7.5 ml blood samples per patient were collected into CellSave® tubes (Menarini Silicon Biosystems, Italy). The CellSearch® Epithelial Cell Test (Menarini Silicon Biosystems, Italy) was applied for CTC enrichment and enumeration according to the instruction from the manufacturer. Samples from the SUCCESS study were prepared using a slightly modified protocol, pooling three separate CellSave® tubes (30 ml) as described elsewhere ^52^. CTC-positive cartridges were sent from clinical centers to the Chair of Experimental Medicine and Therapy Research, Regensburg for cell isolation and molecular analysis. Cells were extracted from CellSearch^®^ cartridges and isolated using the DEPArray^TM^ system (Menarini Silicon Biosystems, Italy) and single cell DNA was amplified by whole genome amplification for subsequent *PIK3CA* mutation analysis.

### Isolation of human primary mammary epithelial cells

Primary human non-cancerous mammary tissue was dissociated as previously described ^20^. Briefly, upon mechanical digestion the tissue was subjected to enzymatic digestion overnight at 37°C in DMEM/F12 (Pan-Biotech, Germany) supplemented with 10mM HEPES (Sigma-Aldrich, Germany), 2% bovine serum albumin (Sigma-Aldrich, Germany), 5 μg/ml insulin, 0.5 μg/ml hydrocortisone, 10 ng/ml cholera toxin (Sigma-Aldrich, Germany), 300 Units/ml collagenase and 100 Units/ml hyaluronidase (all from Sigma-Aldrich, Germany). After removal of organoids and adipocytes by centrifugation at 210 g for 2 min, the cell suspension was passed over a 100 μm and 40 μm cell strainer to obtain a single cell suspension. Separation of fibroblasts from epithelial cells was accomplished by centrifugation at 350 g for 4 min and epithelial cells from the cell pellet were cultured as mammospheres.

### Mammosphere culture

Cell lines and primary HMECs were seeded at a density of 10,000 cells/ml and 50,000 cells/ml, respectively, in 3 cm, 6 cm or 10 cm cell culture dishes or 96 well flat-bottom plates (Thermo Fisher Scientific, Germany; Sigma-Aldrich, Germany; TPP AG, Switzerland). For analyses using the Operetta high content imaging system cells were plated with 10,000-50,000 cells/ml in 96-well µClear plates (Greiner Bio-One, Germany). To prevent attachment of cells all dishes/plates were coated with polyhydroxyethylmethacrylate (PolyHEMA) (12 mg/ml in 95% ethanol, Sigma-Aldrich, Germany) overnight. PolyHEMA-coated dishes/plates were UV-sterilized for 30 min. Cells were cultured in mammosphere medium consisting of MEBM (Lonza, Germany) supplemented with 1% penicillin/streptomycin (Sigma-Aldrich, Germany), 1xB27 (Life Technologies, Germany), 10 ng/ml EGF (Sigma-Aldrich, Germany), 10 ng/ml bFGF (Sigma-Aldrich, Germany), 4 µg/ml heparin (Sigma-Aldrich, Germany) and 1% methylcellulose, if the Operetta-high content imaging system was used. For some analyses mammosphere media was supplemented additionally with 10 ng/ml IL6 (Sigma-Aldrich, Germany), 1.5 µg/ml anti-IL6 antibody (Sigma-Aldrich, Germany), 20 ng/ml Hyper-IL6, 0.1 or 10 ng/ml recombinant human sgp130-Fc (R&D Systems, Germany). Hyper-IL6 was a kind gift of Stefan Rose-John, Christian-Albrechts-University, Germany. Mammospheres were cultured in a humidified atmosphere with 5.5% CO2 and 7% O2 at 37°C for 4 or 7 days.

For setting-up of secondary mammosphere cultures, conducting flowcytometric or single cell expression analyses, first generation mammospheres were collected on day 7 by gentle centrifugation (100 g), dissociated into single cell suspension with trypsin-EDTA (Pan-Biotech, Germany) for 3 min followed by trypsin neutralizing solution (Lonza, Germany). Single cell suspensions of secondary mammospheres were obtained as described for day 7-first generation mammospheres.

### Mammosphere counting

The number of spheres with a diameter ≥ 50 µm was determined by manually counting of a complete plate/dish at day 7 using an inverted microscope (Olympus, 10xair objective). Alternatively, spheres were counted using the Operetta CLS high-content imaging system (PerkinElmer, Hamburg, Germany) by adding CyTRAK Orange (BioStatus Ltd, United Kingdom) at day 4 to the wells at a final concentration of 10 µM. After 60 min incubation, fluorescence imaging of the plates was performed using a 5x air objective and imaging of nine regions per well that were stitched to cover the entire well surface. Harmony high content analysis software was used to analyze the images and to count formation of spheres with diameter ≥ 50 µm (Version 4.8; PerkinElmer, Hamburg, Germany).

### Isolation of LRCs, nLRCs, QSCs from mammosphere cultures

HMECs were labeled with the PKH26 red fluorescent cell linker kit (Sigma-Aldrich, Germany) at 40 nM for 2 min at RT, the reaction was stopped with 10% FBS containing medium and cells were washed three times before plating into primary or secondary mammosphere cultures. LRCs and nLRCs were isolated from spheres of secondary mammosphere cultures that were dissociated with trypsin-EDTA (Pan-Biotech, Germany) for 3 min, neutralized with trypsin neutralizing solution (Lonza, Germany) and stained with DAPI (Roche Diagnostics, Germany) for live/dead cell discrimination. Single LRCs and nLRCs were isolated as single PKH-positive/DAPI-negative and PKH-negative/DAPI-negative cells using a micromanipulator (Eppendorf PatchMan NP2, Eppendorf, Germany) or flow cytometric activated cell sorting. QSCs were isolated at day 7 from primary mammosphere cultures as single, DAPI-negative/EpCAM-positive/PKH-positive cells that did not form spheres and using the micromanipulator.

For flow cytometric assessment of proliferation of MCF 10A-LRCs and MCF 10A-nLRCs, MCF 10A cells were labeled with CFDA-SE (ebioscience, Germany) at 2 µM for 10 min at 37°C in PBS/1% FBS, washed twice after stopping of the reaction with 10% FBS containing medium and cultured as mammospheres. At day 4, single cell suspensions were obtained as described above and analyzed by flowcytometry.

### In vitro differentiation of HMECs

matrigel (growth factor reduced, without phenol-red, BD Biosciences, Germany) was diluted 1:1 with differentiation medium (Ham’s Dulbecco’s modified Eagle’s/F12 medium (Pan-Biotech, Germany), 5% fetal calf serum (Pan-Biotech, Germany), 5 μg/mL insulin (Sigma-Aldrich, Germany), 1 μg/mL hydrocortisone (Sigma-Aldrich, Germany), 10 μg/mL cholera toxin (Sigma-Aldrich, Germany), 10 ng/mL EGF (Sigma-Aldrich, Germany), 1× penicillin/streptomycin/fungizone, (Lonza, Germany)), smeared in 2 well slides and incubated for 15 min. at 37°C. On top, 50,000 cells from disaggregated secondary mammospheres were added. After incubation for 30 min at 37°C cells were covered with an additional matrigel layer and incubated for additional 15 minutes at 37 ⁰C. Differentiation medium was added at the end of the embedding procedure and exchanged every two days. Cultures were examined 3-4 weeks post embedding for the development of tubular and acinar structures.

### Culture of primary human mesenchymal stem cells and generation of osteoblasts

Mononuclear cells from bone marrow aspirates were cultured at a density of 2×10^6^ cells in a T75 flask (Sarstedt, Germany) in DMEM with 1g/L glucose, 4mM glutamine and 1mM sodium pyruvate (all from Life Technologies, Germany), supplemented with 10% MSC-qualified FBS (WKS Diagnostik, Germany), 1% penicillin/streptomycin (Life Technologies, Germany) and 1ng/ml bFGF (Peprotech, Germany). Adherent cells were cultured for 3 weeks and cryo-conserved. Before cryo-conservation MSCs were tested for the expression of CD45, CD34, CD90, CD105, CD44 and Nestin by flow cytometry (Supplementary Fig. 5). Also, the ability of MSCs to differentiate into adipocytes and osteoblasts was tested as previously described ^71^. Briefly, osteoblasts for co-culture experiments were generated by culturing confluent MSC-cultures in DMEM with high glucose (Life Technologies, Germany) supplemented with 10% MSC-qualified FBS (WKS Diagnostik, Frankfurt, Germany) 1% penicillin/streptomycin (Life Technologies, Germany), 10^-7^ M Dexamethasone, 25 µg/ml L-ascorbic acid and 3 mM sodium dihydrogen phosphate (all from Sigma-Aldrich, Germany) for 21 days with medium being changed every other day.

### Co-cultures of MCF10A with MSCs, OBs, HUVECs

MSCs and HUVECs were plated at a density of 4×10^5^ cells/well of a 6 well plate (Corning, Germany) in their respective growth medium. The next day, medium was exchanged to MCF 10A growth medium and 1×10^5^ MCF 10A-GFP cells were added to each well. In case of co-cultures with OBs, 4×10^5^ MSCs per 9.6 cm^2^ surface of a 6 well plate (Corning, Germany) were plated and differentiated into OBs for 21 days. On day 22 medium was exchanged to MCF 10A growth medium and 1×10^5^ MCF 10A-GFP cells were added to each well. For cultures with transwells, MSCs were plated at 1.75×10^5^ cells per 4.2cm^2^ of a 6-well transwell insert (Falcon 353090, VWR, Germany).

### Xenotransplantations of DCCs, HMECs and MDA-MB-231 cells

For xenotransplantations of DCCs, mononuclear cells from BM-aspirates of non-metastasized or metastasized breast or prostate cancer patients were enriched for human EpCAM or depleted of human CD45^+^CD33^+^CD11b^+^cells and erythrocytes using a mix of CD45, CD33, CD11b and Glycophorin A microbeads according to the manufacturer’s instructions (Miltenyi Biotec, Germany). Each sample was then split into halves: one half was subjected to DCC-enumeration by staining for CK8/18/19 or EpCAM. The other half of the cell suspension was transplanted without *ex vivo* expansion into NSG-mice using one to two injection routes (non-metastasized patients) or three to four different injection routes (metastasized patients). In some cases mononuclear cells were cultured as mammospheres in 6 cm culture plates coated with polyhydroxyethylmethacrylate (12 mg/ml, Sigma), under hypoxic conditions (7% O2) at 37°C and in mammosphere medium containing 10 nM HEPES (Sigma-Aldrich, Germany), 10 µg/ml insulin (all from PAN-Biotech, Germany), 5 ng/ml GRO-α (R&D Systems, Germany), 20 ng/ml hyper interleukin-6 (kindly provided by S. Rose-John) and 0.2% Methylcellulose (Sigma-Aldrich, Germany). Cultures were monitored weekly for sphere growth.

To transplant spheres or EpCAM-enriched or CD45/CD11b/erythrocyte depleted bone marrow, cells/spheres were collected in a microwell (volume 10-15 µl, Terasaki, Greiner Bio-One, Germany) pre-coated with polyhydroxyethylmethacrylate (12 mg/mL, Sigma-Aldrich, Germany). Cells or spheres were transplanted in a final volume of 30 µl and 25% high-concentration matrigel (BD Biosciences, Germany) as published before ^9^. Cells were injected with an insulin syringe (Microfine, 29G, U-50, BD Biosciences, Germany) sub-cutaneously, intra-venously, intra-femorally or sub-renally in 4-8 weeks old male or female NSG or NOD.Cg-Prkdc^scid^ mice. Mammary fat pad injections were performed in the 4^th^ pre-cleared mammary fat pad of 3 weeks old female mice in 50% matrigel (BD Biosciences, Germany). Breast or prostate cancer-origin of xenografts were verified by a pathologist.

To assess the differentiation ability of HMEC-spheres *in vivo*, secondary mammospheres were dissociated and 200,000 cells were mixed with 225,000 pre-irradiated (15 Gy) C3H10T1/2 mouse fibroblasts. The cell suspension was then mixed 1:1 with matrigel (growth factor reduced without phenolred, BD Bioscience, Germany) and injected in the 4^th^ pre-cleared mammary fat pad of 3 weeks old female NSG mice. Mice were euthanized 8 weeks after transplantation and analyzed for the presence of human mammary gland tissue.

MDA-MB-231 cells grown under adherent conditions were pre-treated with PBS, an anti-IL6 antibody (1.5 µg/ml, Sigma-Aldrich, Germany) or HIL6 (20 ng/ml, kind gift of Stefan Rose-John, Christian-Albrechts-University, Germany) for 3 hours and 20,000 cells were transplanted into the mammary fat pad of NSG mice as 1:1 mixture with matrigel (BD Biosciences, Germany) in the 4^th^ pre-cleared mammary fat pad of 3 weeks old female NSG-mice. All mice were analyzed when first tumors reached a diameter of about 10 mm.

### Detection of human DCCs and mammary gland in NSG-mice

Lung and bone marrow of mice transplanted with human DCCs were analyzed for the presence of DCCs or metastasis. Lungs were examined by a pathologist. For identification of disseminated cancer cells in the mouse bone marrow, mononuclear cells were screened accordingly to the method for human DCCs using immunofluorescent staining of the cell suspension with anti-human EpCAM (Ber-EP4-FITC, Agilent, or HEA-125-PE, Miltenyi-Biotec Germany) or adhesive slides and staining anti cytokeratin 8/18/19 (A45-B/B3, AS Diagnostik, Germany). Unspecific binding was blocked by using PBS with 5% human AB serum (Bio-Rad, Germany) and 5% mouse serum (Agilent, Germany). Positive cells were isolated with the micromanipulator and subjected to whole genome amplification.

Mammary glands of mice transplanted with HMECs were dissected, fixed and stained with an anti-human cytokeratin 18 antibody (20 µg/ml, clone CK2, Millipore, Germany). Cells expressing human cytokeratin 18 were laser-microdissected (PALM Microbeam system, Bernried, Germany) and subjected to whole genome amplification.

The human origin of DCCs or CK18^+^ cells isolated from mouse bone marrow or laser-microdissected from mammary glands of NSG-mice was confirmed by a PCR discriminating between the human and mouse cytokeratin 19 gene: forward primer: 5’-TTC ATG CTC AGC TGT GAC TG-3’ and reverse primer 5’-GAA GAT CCG CGA CTG GTA C-3’, annealing 58°C, amplicon 621 bp for the human sequence.

### Quantification of HER2 and PGR staining in tissue sections by TissueFAX cytometry

Tissue sections were stained with an automated staining machine (Ventana: BenchMark ULTRA). Tissue sections used for analysis were stained within the same run. Images of stained tissue sections were scanned with the TissueFAXSi-plus imaging system (TissueGnostics, Vienna, Austria; acquisition software: TissueFAXS version 3.5.129) equipped with a digital Pixelink colour camera (PCO AG, Kehlheim, Germany). Images for the analysis of Ki-67 staining were analysed with HistoQuest software version 6.0.1.130 (TissueGnostics, Vienna, Austria). Using that software, two markers were created: hematoxylin as ‘master marker’ (nucleus) and Ki-67 as ‘non-master marker’. To achieve optimal cell detection, the following parameters were adjusted: (i) nuclei size; (ii) discrimination by area; (iii) discrimination by gray and (iv) background threshold. For the evaluation of the percentage of Ki-67expressing cells, scatter plots were created, allowing the visualization of corresponding cells in the source region of interest using the real-time back gating feature. The cut-off discriminated between false events and specific signals according to cell size and intensity of Ki-67 staining. For the PBS-group 112,098 cells (33 regions, 8.25 mm^2^), anti-IL6 group 161,279 cells (41 regions, 10.25 mm^2^) and HIL6-group 98,812 cells (26 regions, 6.50 mm^2^) were analysed.

### Whole genome amplification and analysis of copy number alterations

Single-cell genomic DNA was subjected to whole genome amplification (WGA) using the previously described ^72, 73^ or the commercially available version (*Ampli1*^TM^ WGA, Menarini Silicon Biosystems). DCCs isolated from BM of patients or NSG-mice were subjected to CNA analysis as previously described (mCGH) ^72, 73^ or using the *Ampli1*^TM^ LowPass kit (Menarini Silicon Biosystems) according to the manufacturer’s instructions.

### *PIK3CA* sequencing of single cells

*PIK3CA* mutation in CTCs and DCCs was assessed using the *Ampli*1^TM^ PIK3CA Seq kit (Menarini Silicon Biosystems) or amplicon-based sequencing on single cells following *Ampli* 1^TM^ WGA. For amplicon-based sequencing the following primers were used: Exon 9 forward primer 5’-AAG CAA TTT CTA CAC GAG A-3’ and reverse primer 5’-CC TTA TTT ATT TCG TCT TAA ATG-3’, annealing 58°C, amplicon size 189 bp; Exon 20 forward primer 5’-TCT AGC TAT TCG ACA GCA TGC −3’ and reverse primer 5’-T ACC TAA CCT AGA AGG TGT GTT −3’ annealing 58°C, amplicon size 221 bp. For each exon, 1 µl of WGA-product of CTCs or DCCs was used for the PCR. Resulting products were loaded on a 1.5% agarose gel and negative PCR results were considered dropouts for *PIK3CA* analysis. Positive CTC samples were purified using QIAquick purification kit (Qiagen, Germany) according to the manufacturer’s protocol with the exception that elution at the end of the protocol was in 25 µl water. Purified CTC samples were sent to a sequencing provider (Sequiserve, Germany). PCR products from positive DCC samples were purified by the sequencing provider (GATC, Germany).

### Whole transcriptome amplification (WTA) of single spheres and cells

Whole transcriptome amplification of single cells or undissociated spheres was performed as previously described ^19, 74^. The quality of WTA products was assessed by expression analysis of three housekeeping genes: *EEF1A1*, *ACTB* and *GAPDH*. Only samples positive for all three markers were used for downstream analyses.

### mRNA microarray experiments

MCF 10A cells were cultured as mammospheres in the presence or absence of 10 ng/ml IL6 (Sigma-Aldrich, Germany), 10 ng/ml IL6 + 0.1 ng/ml recombinant human sgp130-Fc (R&D Systems, Germany) or 20 ng/ml Hyper-IL6 (kind gift of S. Rose-John, Christian-Albrechts-University, Germany) for 12 and 24 hours. Cells were seed in triplicates for all conditions and time points. After 12 and 24 hours, cells were collected by centrifugation (5 min at 500 x g) and RNA was isolated using RNeasy Mini Kit (RNeasy Mini Kit, Qiagen, Germany) according to the manufacturer’s protocol. Microarray analysis was performed using the Whole Human Genome Microarray Kit, 4×44K (G4112F, Agilent Technologies, Germany).

For transcriptome analysis of LRCs, nLRCs and QSCs, HMECs were cultured and cells isolated as described above and cDNA was obtained from manually isolated single cells using whole transcriptome amplification.

Labelling of cDNA was performed by PCR with Cy5-labelled primers. Reaction mix contained 5 μl of buffer I (Expand Long Template, Roche, Germany), 3% (v/v) deionized formamide, 0.35 mM each dNTP, 2.5 μM 5’-U*CAGAAU*TCAUGCCC*CCCC*CCCC*C-3’ primer (*denotes nucleotides conjugated with Cy5 fluorophore; Metabion), 3.75 U of PolMix (Expand Long Template, Roche, Germany) and 1 μl of WTA-product or 100 ng cDNA from bulk RNA preparations of MCF 10A cells in a final volume of 49 μl. PCR parameters were: one cycle with 1 min at 95 °C, 11 cycles with 15 s at 94 °C, 1 min at 60 °C, and 3 min 30 s at 65 °C, 3 cycles where the elongation time was increased 10 s per cycle, and finally one cycle with an elongation time of 7 min. Labelled products were purified using a PCR purification kit (Qiagen, Germany) according to the instructions of the vendor. Purified Cy5-labelled DNA was denatured by incubation for 5 min at 95 °C followed by incubation on ice. Hybridization solution was prepared by mixing 42 μl of denatured Cy5-labelled DNA, 55 μl of 2x HiRPM hybridization buffer (Agilent, Germany), 11 μl of 10X GE Blocking agent (Agilent, Germany), 4 μl of 25% (v/v) Tween-20, and 4 μl of 25% (v/v) Igepal. Four 100 μl samples of hybridization mix were overlaid on four hybridization fields of Agilent Whole Human Genome (4×44K) Microarray Kit with SurePrint microarray slides and incubated for 17 h at 65 °C under constant rotation. After hybridization, slides were washed in Agilent Wash buffer 1 for 1 min on a shaker in the dark and incubation continued in Agilent Wash buffer 2 pre-warmed to 37 °C. Slides were dried by washing for 30 s in acetonitrile and scanned on a GenePix 4400 A scanner (Molecular Devices, Germany). Numerical readouts of fluorescence intensities (GPR files) were generated using GenePixPro 7 (Molecular Devices, Germany).

### NGS mRNA library preparation and sequencing

The majority of the (TTT)7 and (CCC)5 nucleotides forming the ends of cDNA products were removed by a limited-cycle PCR with primers introducing BpuEI and BglII restriction sites followed by restriction enzyme digestion. Briefly, 1µl of a 1/5 dilution of the original WTA sample was used in a total volume of 20µl with 24µM of primer CP2-BpuEI (5’-TCA GAA TTC ATG (CCC)5 GTC TTG AGT TTT TT-3’) and 24 µM of primer Cp2-BglI-13C (5’-TCA GAA TTC ATG (CCC)2 CGG (CCC)2-3’) for amplification. After an initial denaturation at 95°C for 1min, 5 cycles of 94°C for 15 sec, 60°C for 1 min, and 65°C for 210 sec and 3 cycles of 94°C for 15 sec, 60°C for 1min, and 65°C for 210 sec (+10 sec/cycle) were carried out followed by a final extension step of 7 min. Resulting cDNA products were purified with 1.8 volume of Ampure XP beads (Beckman Coulter, USA) according to the manufacturer’s instructions and eluted in 40 µl of distilled water. Five µl of EcoRI buffer supplemented with 80 µM S-adenosyl methionine (New England Biolabs, Germany) and 2.5 µl BpuEI (5 U/µl) were added in a volume of 50 µl and incubated at 37°C for 1 hr followed by heat inactivation of the enzyme for 20 min at 65°C. Subsequently 1 µl of EcoRI buffer supplemented with 80 µM S-adenosyl methionine and 2.5µl BglII (10 U/µl) were added in a final volume of 60 µl and incubated 3 hrs at 37°C followed by heat inactivation of the enzyme. The complete restriction digest was purified with 1.8 volume of Ampure XP beads according to the manufacturer’s instructions and eluted in 16 µl 10 mM Tris-Cl, pH 8.5 (Elution buffer EB, Qiagen, Germany). The length distribution of purified cDNA populations was determined on the Bioanalyzer 2100 (Agilent Technologies, USA). Optimal Covaris settings for fragmentation of each purified cDNA sample to 350 bp insert size were determined on the basis of the average length distribution. Subsequently sequencing libraries were prepared according to the TruSeq DNA PCR-Free Library Prep Kit (Illumina, USA). Resulting libraries were quantified with KAPA Library Quantification Kit for Illumina Platforms (Kapa Biosystems, RSA), pooled in equal molar ratios and sequenced on Illumina NovaSeq 6000 platforms.

### IL6, IL6RA and gp130 mRNA expression analysis in single cells

IL6, membrane IL6 receptor, spliced IL6 receptor and gp130 expression was assessed by PCR using the MJ Research Peltier Thermal Cycler Tetrad (Bio-Rad, Germany) with the following primers: IL6 (forward primer 5’-GAG AAA GGA GAC ATG TAA CAA GAG T -3’, reverse primer 5’-GCG CAG AAT GAG ATG AGT TGT -3’, annealing 62°C, amplicon size 388 bp), membrane versus spliced IL6RA (forward primer 5’-CTG CAA ATG CGA CAA GCC TC -3’, reverse primer 5’-GTG CCA CCC AGC CAG CTA TC - 3’, annealing 62°C). The spliced and membrane-bound IL6 receptor can be distinguished according to their PCR product size: mIL6RA 380 bp, spliced IL6RA 286 bp. Gp130 forward primer was 5’-GGA CCA AAG ATG CCT CAA CT -3’, reverse primer 5’-GGC AAT GTC TTC CAC ACG A -3’, annealing 58°C and amplicon size 280 bp. gp130 expression was assessed by quantitative PCR on re-amplified and purified (Qiagen PCR Purification Kit, Qiagen, Germany) WTA products of single cells as previously described ^74^. To normalize for the template input quantification of yields in the individual samples was spectrophotometrically conducted using the NanoDrop 2000 instrument. The DNA input for each qPCR of a single cell was normalized to 2.5 ng and the qPCR run as previously described with the following primers: *gp130* forward primer 5’-ATA TTG CCC AGT GGT CAC CT -3’ and reverse 5’-AGG CTT TTT GTC ATT TGC TTC T -3’, annealing 58°C, amplicon size 125 bp. Fold changes in gp130 expression were calculated from the delta Cp-values between MCF 10A or MCF-7 cultured with and without MSCs.

### Flowcytometry

Spheres or adherent cells were trypsinized with trypsin/EDTA (Pan-Biotech, Germany) for 3 min, if not stated otherwise. MSC monocultures and co-cultures of MCF10A-GFP cells with MSCs, OBs and HUVECs were harvested by trypsin/EDTA (Pan-Biotech, Germany) for 5 min and using cell-scrapers. To reduce non-specific binding single cell suspensions were incubated for 5 min at 4°C with PBS/10% AB-serum (Bio-Rad, Germany), subsequently stained with fluorescence-labeled or biotinylated antibodies for 15 min at 4°C and washed once with PBS/2% FCS/0.01% NaN3. In case of biotinylated primary antibodies, PE-labeled streptavidin (Dianova, Germany) was used as secondary staining reagent. Cells were stained using the following antibodies: anti-human CD24-APC (ML5), anti-human CD34-PE (581), anti-human CD44-V450 (G44-26), anti-human CD45-FITC, APC or PerCP-Cy5.5 (HI30), anti-human CD90 Alexa Flour 700 (5E10), anti-human CD105-FITC (43A3), anti-human CD130-APC (2E1B02), anti-human Nestin-PE (10C2), biotinylated anti-human IL6R (UV4), isotype control mouse IgG2a-APC (MOPC-21), isotype control mouse IgG2b-V450 (MOPC-21), isotype control IgG1-biotin (MOPC-21) (all purchased from BioLegend, Germany) and anti-human EpCAM (HEA-125, Miltenyi-Biotech, Germany). Viability dye eFlour 780 (ebioscience, Germany) was used for live/dead cell discrimination. Cells were analyzed on a LSR II machine equipped with FACS DIVA 5.03 software (BD Bioscience, Germany) and data was analyzed with FloJo 8.8.6, 10.1 or 10.5.3 (Treestar, USA). Sorting of PKH26-labeled LRC and nLRCs was performed with a FACSAria cell sorter (BD Bioscience, Germany).

### IL6 and soluble IL6RA detection by ELISA

IL6 and soluble IL6RA concentrations were assessed in 100 µl cultured media obtained from HMECs or MCF 10A cells propagated under anchorage dependent or anchorage independent conditions with the *Human IL-6 DuoSet* or *Human sIL-6R alpha DuoSet* ELISA kit (R&D Systems, Germany) following the manufacturer’s recommendations.

### Inhibition of ADAM-proteases

MCF 10A cells were treated for 48 hrs with 20 µM TAPI-2 acetate salt (Sigma-Aldrich, Germany). The culture supernatant was tested for the presence of IL6 and sIL6RA by ELISA.

### Immuno-(western) blotting

Cell lysates were prepared using ice cold RIPA Buffer supplemented with cOmplete, EDTA-free Protease Inhibitor Cocktail and PhosSTOP^TM^ (all from Sigma-Aldrich, USA). The Protein concentration of lysates was determined with Pierce^TM^ BCA Protein Assay Kit (Thermo Fisher Scientific, USA). Cell lysates were mixed with 4x Laemmli Sample Buffer (Bio-Rad, USA) containing 10% 2-Mercaptoethanol (Sigma-Aldrich, USA) and denatured for 5 min at 95°C. 10 µg of protein/lane were loaded on 12% Mini-PROTEAN® TGX^TM^ Gels (Bio-Rad, USA) and protein separation was performed with SDS PAGE Running Buffer (25 mM Tris, 192 mM glycine, 0,1% SDS). Proteins were blotted onto Immobilon-P PVDF Membranes (Millipore, USA). For washing of membranes TBS-T (137 mM NaCl, 20 mM Tris, 0,05% *(w/v)* Tween-20, pH 7.6) was used. To detect signaling protein, the following primary antibodies (all from Cell Signaling Technology, USA) were used at dilutions according to manufacturer’ s instructions: anti-phospho-STAT3^Tyr705^ (clone D3A7), anti-phospho-AKT^Ser473^ (clone D9E), anti-phospho-ERK1/2^Thr202/Tyr204^ (clone E10), anti-STAT3 (clone 124H6), anti-AKT (clone 40D4) and anti-ERK1/2 (clone 137F5). As loading control an anti-α-Tubulin antibody (Sigma-Aldrich, USA, clone DM1A; 1:5000) was used. This was followed by incubation with horseradish peroxidase (HRP)-conjugated goat anti-rabbit IgGs or goat anti-mouse IgGs (both Sigma-Aldrich, USA; 1:10000). Protein bands were visualized using SuperSignal^TM^ West Pico PLUS Chemiluminescent Substrate (Thermo Fisher Scientific, USA). Chemiluminescence was recorded by a ChemiDoc^TM^ MP Imaging System and analyzed with Image Lab^TM^ Software (both Bio-Rad, USA). Membranes were stripped for re-probing using Restore ™ Plus Western Blot Stripping Buffer (Thermo Fisher Scientific, USA) according to manufacturer’s instructions.

### Bioinformatics

#### MCF 10A HIL6/IL6-stimmulation mRNA microarray data

Gene expression data were obtained using the Agilent Whole Human Genome Microarray Kit (4×44K) and quality assessed by inspection of chip raw images and gene expression frequency distributions. All 24 expression profiles (3 biological replicates, 4 treatment groups, 2 time points) were of sufficiently high quality for further bioinformatic analysis. Raw gene expression data were background corrected (limma Bioconductor-package ^75^, version 3.36.5, normexp method), log2-transformed and normalized by quantile normalization. Replicated probes (identical Agilent IDs) were replaced by their median per sample. Gene ranking was performed using empirical array quality weights ^76^ and linear models from the limma Bioconductor-package (version 3.36.5) using standard treatment versus control contrasts. Gene annotation (Supplementary Table 3) was obtained by aligning Agilent oligo sequences to NCBI RefSeq genes (August 8, 2019) using BLAST ^77^ (version 2.9.0) requiring 100% identical matches, a maximum length difference between oligo and target sequence of one, and less than 100 hits per oligo. In addition, ensembl annotation ^78^ (version 97) was retrieved and used as a secondary information source (e.g. for oligos that were unannotated by NCBI RefSeq). GENCODE metadata (version 32) ^79^ were used as complementary annotation. For gene lists, graphical display and functional annotation, probes targeting the same gene were disambiguated by retaining only the probe with the lowest *P value*. Differential gene expression was defined by a maximum FDR-adjusted *P value* of 0.05 and a minimum absolute log2-fold change of log2(1.5) = 0.58. Computations were performed using R version 3.5.1 ^80^.

#### Mammary cell subpopulation mRNA microarray data

Human mRNA expression data from Lim et al. ^41^ based on Illumina HumanWG-6 v3.0 BeadChip microarrays were downloaded from the Gene Expression Omnibus (GEO) (series GSE16997). Data pre-processing, analysis and annotation was performed analogous to the procedure detailed above for MCF 10A cells except that the linear model included all pairwise contrasts between the three cell types.

Fold change analysis of the MCF 10A and mammary cell subpopulation data was performed by first selecting a pairwise comparison from the MCF 10A data (e.g. classical IL6 stimulation vs. control) and another from the mammary subpopulation data (e.g. luminal progenitor vs. mature luminal), each performed according to moderated t-testing (limma Bioconductor-package, version 3.36.5). The differential gene lists of both comparisons were intersected and the randomness of their overlap quantified using hypergeometric testing (Supplementary Table 6). Second, the log-fold-changes of both comparisons were correlated without centering (i.e. without subtracting the respective group means) because reference to zero log-fold was intended. Correlation *P values* were calculated according to centered Pearson correlation.

#### LRC/QSC/nLRC mRNA microarray data

Gene expression data were obtained using the Agilent Whole Human Genome Microarray Kit (4×44K). All chips passed quality assessment and were pre-processed and annotated as described for MCF 10A cells above except that no fold-change limit (originally used in the data of Lim et al. ^41^ and thus also employed for MCF 10A cells) was applied. The data showed patient effects that were accounted for by including patient IDs as second covariate in the linear model after safeguarding independence between patients and sample groups: Cramer’s V with bias correction = 0; R-package rcompanion version 2.3.7 ^81^. For graphical display these effects were compensated by using the function removeBatchEffect from the Bioconductor package limma (version 3.36.5). Dimension reduction to 2D according to t-SNE ^82^ (Fig. 2c) and pairwise differential expression analysis (number of differentially expressed genes was: 35 for nLRC vs. QSC, 127 for LRC vs. nLRC and 163 for LRC vs QSC; FDR-adjusted P value < 0.05, Supplementary Table 2) revealed that nLRC and QSC were much more similar to each other as compared to LRC. To concentrate on main effects, nLRC and QSC were pooled resulting in 216 differentially expressed genes for LRC vs (nLRC+QSC). Enrichment analysis was aimed at the NCI-Nature Pathway Interaction Database ^83^ for its focus on cancer research and treatment and conducted using the R-package enrichR ^84^ (version 2.1).

#### DCC and HD mRNA sequencing data

The sequencing quality was evaluated per sample with FastQC ^85^ and in a multi-sample comparison with MultiQC ^86^ and the independent tool MusaQC before and after adapter trimming and contamination screening. Briefly, raw sequencing data of single cells (30 M0-, 11 M1-stage DCCs and 15 EpCAM+ cells from healthy donors (HD) from 21, 5 and 7 patients, respectively) were trimmed and remaining adapter sequences as well as low sequencing quality bases at the end of each read were removed using BBDuk ^87^. In order to increase the mapping quality (lowering false positive alignments), read decontamination was performed using BioBloom Tools ^88^ with filters for the genomes of *Homo sapiens* (hg38), *Mus musculus* (mm10), *Escherichia coli* (BL21), *Mycoplasma pneumoniae* (M129), *Sphingobium* sp (SYK-6), *Bradyrhizobium japonicum* (USDA 110), *Pichia pastoris* (GS115), *Malessia globosa* (CBS 7966), *Aspergillus fumigatus* (Af293) and a set of viral genomes (RefSeq, 5k+ genomes). All reads that did not map exclusively to hg38 (GENCODE version 27, GRCh38.p10) or did not map at all were defined as likely contaminations and discarded from downstream processing. Subsequently, the cleaned sample reads were aligned to the reference genome hg38 with STAR (version 2.5.1b) ^89^. Uniquely mapped reads were counted per gene per sample using featureCounts from Subread ^90^. We performed quality control and checked for outlier samples with the Bioconductor-package scater (version 1.12.2) ^91^ using the functions calculateQCMetrics and plotPCA for QC metrics with outlier detection enabled. The results showed that none of the samples was an outlier. Thus, we kept all samples for further analysis. Samples were sequenced in two batches with only very little association between batches and phenotype (M0/M1/HD): Cramer’s V with bias correction = 0.11; R-package rcompanion^81^ (version 2.3.7). We applied the multiBatchNorm (Bioconductor package: batchelor 1.0.1) to all cells and further rescaleBatches (Bioconductor package: batchelor 1.0.1 ^92^) to DCCs to remove batch effects and get the normalized log2 counts. After batch correction we obtained 8626 and 7359 expressed genes for HD cells and DCCs on average, respectively.

The top 500 most variable genes were analyzed using PCA. From the genes annotated by GO terms containing “B cell”, “Epithelial” or “Epithelium”, the top 100 most variable were subjected to PCA. PCAs were calculated using prcomp (R stats package). For pathway enrichment analysis, we filtered for protein coding genes and compiled 2×2 contingency tables for each sample and each pathway according to whether genes were expressed (log2 (normalized counts)>0) and present in the pathway. Contingency tables were subsequently evaluated according to one-tailed Fisher’s exact test (R stats package). Calculations were performed using R version 3.6.0.

#### Analysis of copy number alterations

To enable the combined analysis of mCGH- and LowPass-Seq-derived CNA profiles, the genomic coordinates obtained with the LowPass bioinformatics analysis pipeline (Menarini Silicon Biosystems, Italy) were converted to cytoband information using a custom script for R ^80^ and the UCSC Goldenpath reference (version hg38 ^93^). Afterwards, the aberrations were manually screened and compared to the respective CNA profile images before being annotated according to the specifications of the International System for Human Cytogenetic Nomenclature (ISCN) ^94^. Small aberrations < 1 megabase as well as recurring technical artefacts in chromosome 1p and centromeric and telomeric regions were excluded. Finally, the combined ISCN-annotated aberration data (mCGH and LowPass-Seq) were stratified into M0 and M1 groups and submitted to the Progenetix user data tool ^95^ to generate individual frequency plots for M0 and M1 cells.

#### Data availability

All genomic results of this study are available within the article and its Supplementary Information.

### Statistical analysis

Statistical analysis was performed using the GraphPad Prism 6.0 software (GraphPad Software, Inc., USA). Differences in mean values between groups were analyzed by Student’s t-test, Mann-Whitney test or one-way ANOVA followed by post-hoc statistical testing, where appropriate. Time dependencies were analyzed by regression analysis (F-test). Independence in contingency tables was assessed by Fisher’s exact test. All tests were realized two-sided. A P value of less than 0.05 was considered statistically significant.

---

**Supplementary Figure 1:**
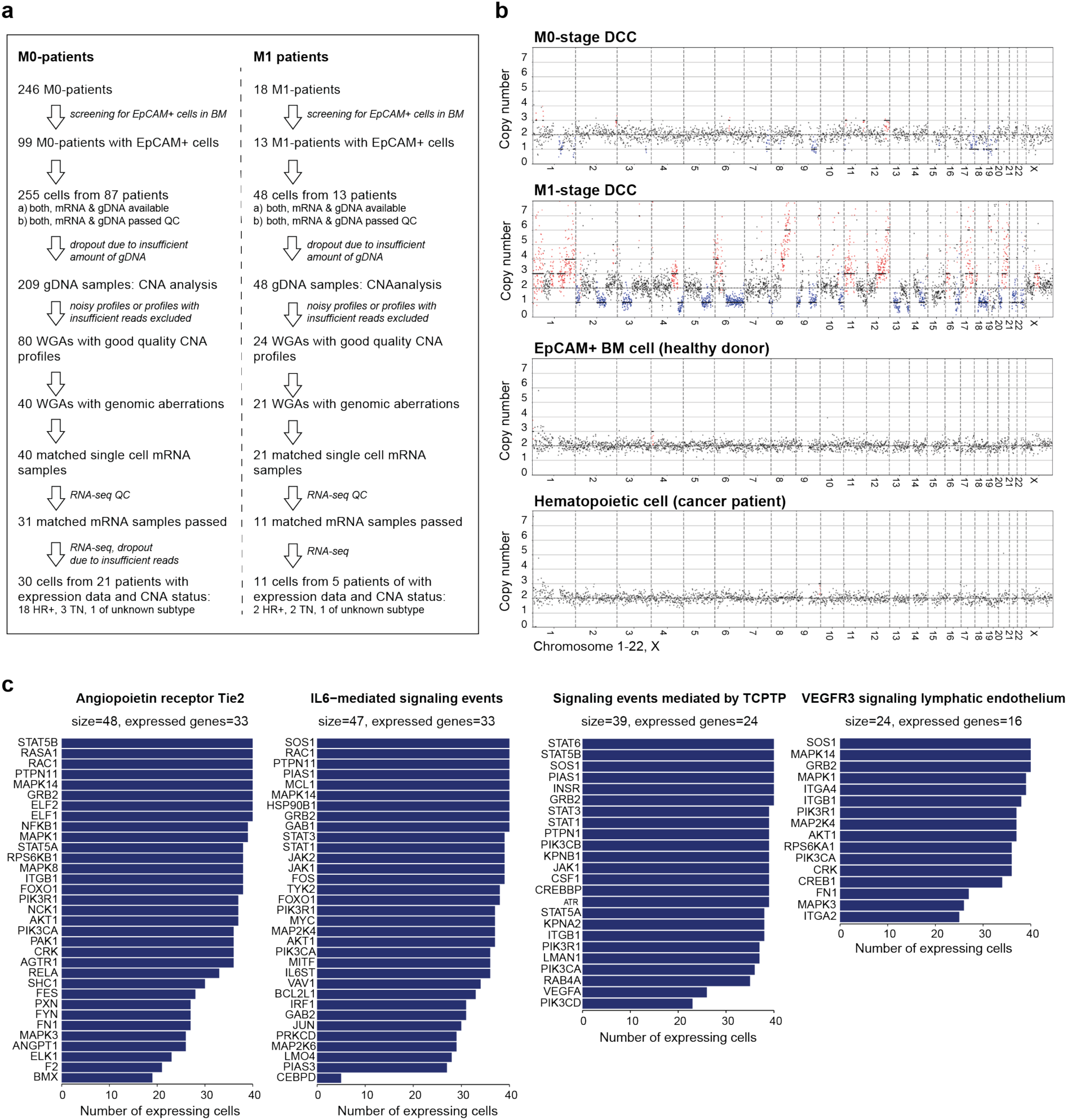
Identification and molecular analysis of DCCs from BM of breast cancer patients. **a** Isolation of EpCAM^+^ DCCs from BM of non-metastasized (M0-stage) and metastasized (M1-stage) breast cancer patients. DNA and RNA were isolated from each cell by WGA and WTA for CNA and RNAseq analysis, respectively. **b** Representative single cell CNA profiles of M0- and M1-stage DCCs and control cells (EpCAM+ cell from BM of a patient without malignant disease or a hematopoietic cell of a cancer patient). **c** Number of DCCs expressing genes of pathways identified to be enriched in DCCs (see Fig. 2c).

**Supplementary Figure 2:**
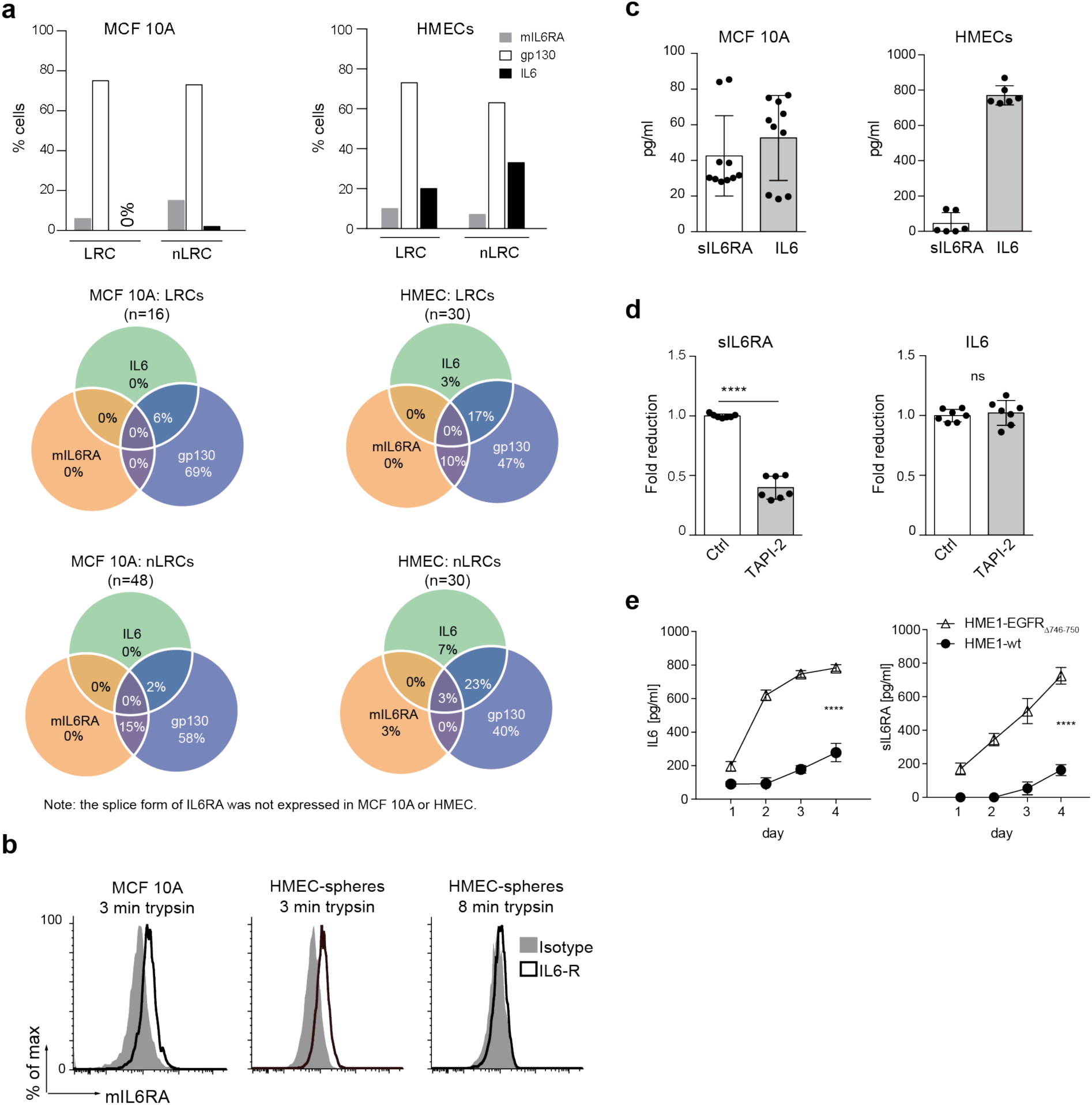
Expression of IL6 signaling molecules in MCF 10A, hTERT-HME1 and primary HMECs. **a** Expression of IL6, mIL6RA and gp130 mRNA in single LRCs or nLRCs from MCF 10A or HMEC-spheres. The spliced soluble form of IL6RA was not found to be expressed. Expression of IL6 signaling molecules did not differ significantly between LRCs and nLRCs of MCF 10A or HMECs (LRCs vs. nLRCs for IL6/mIL6Ra/gp130 in MCF10A or HMECs (Fisher’s exact test, P values for all comparisons >0.05). **b** IL6RA is expressed on the cell surface of MCF 10A cultured under non-sphere conditions and primary HMEC-spheres. The data is representative of three independently performed experiments. **c** IL6 (N=10) and soluble IL6RA (N=10) were measured in the cell culture supernatant of MCF 10A cultured under non-sphere conditions or primary HMEC-spheres (cumulative data of three patients, each patient in duplicate). **d** MCF 10A cells were cultured under non-sphere conditions without (N=7) or with 20 µM TAPI-2 (N=7), an inhibitor of ADAM-proteases. Protein levels of soluble IL6RA (sILRA) and IL6 in the supernatant were determined by ELISA. **e** IL6 and IL6RA in the supernatant of HME1-wt and isogenic HME1-EGFR^Δ746-750^ cells cultured under non-sphere conditions was determined by ELISA. Cumulative data of three experiments, each data point in duplicate. Panel d: two-sided Student’s t-test, panel e: linear regression analysis; asterisks indicate significance * P<0.05, **** P<0.0001. All error bars correspond to standard deviation (Mean ± SD).

**Supplementary Figure 3:**
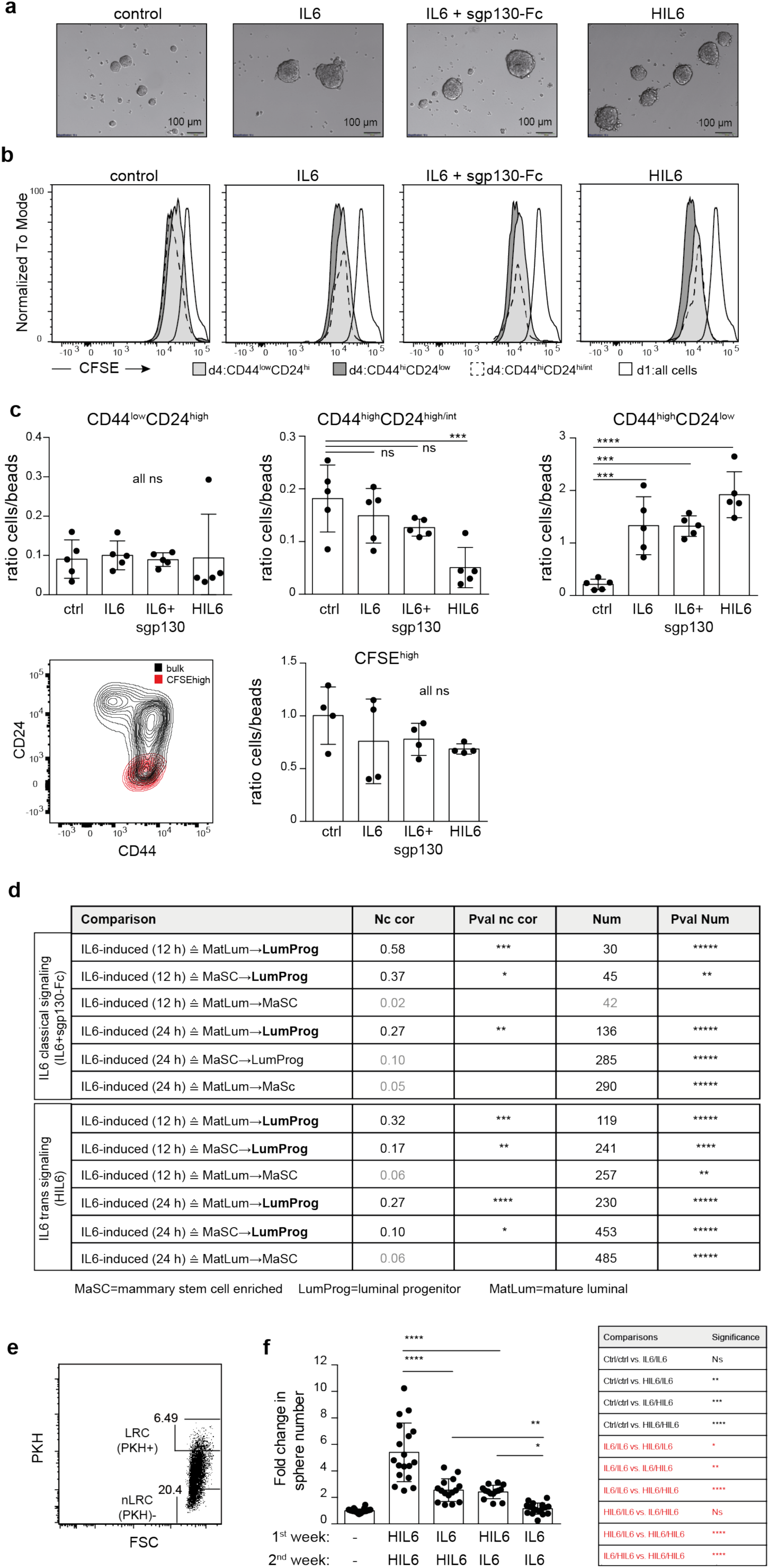
IL6 trans-signaling converts non-stem cells into stem-like cells. **a** MCF 10A spheres cultured without or with IL6, IL6 plus sgp130-Fc or with HIL6. **b** CFSE-labeled MCF 10A cells were cultured as spheres with or without activators (IL6, HIL6) and inhibitors of classical (an anti-IL6 antibody) and trans-signaling (sgp130-Fc). CFSE-dilution in CD44^high^CD24^low^, CD44^high^CD24^high^ and CD44^low^CD24^high/intermediate^ cells was determined by flow cytometry at day 4. The CFSE-fluorescence intensity of all cells at day one is included as reference. Data are representative for three 3 independently performed experiments. **c** The absolute number of CD44^high^CD24^low^, CD44^high^CD24^high^, CD44^low^CD24^high/intermediate^ cells (upper panel) and LRCs (CFSE^high^, lower panel) was determined as cell/bead ratio at day 4 by flow cytometry (N=4-5 per group). **d** Fold-change correlation analysis comparing gene expression changes induced by IL6 plus sgp130 (classical signaling) and HIL6 (trans signaling) in MCF 10 A cells at 12 and 24 hrs with the gene expression signatures of luminal progenitor (LumProg), mature luminal (MatLum) and mammary stem cell enriched cells (MaSC) according to the study of Lim et al. ^35^ Nc cor: non-centered correlation between fold-changes, Num: number of common differentially expressed genes. **e** nLRCs from primary, PKH26-labelled control mammosphere-cultures were sorted by flow cytometry as PKH^-^ cells. **f** Primary HMECs were cultured as spheres for two consecutive rounds in the absence (N=26) or presence of HIL6 and IL6 (HIL6+HIL6, N=18; IL6+HIL6, N=15; HIL6+IL6, N=14; IL6+IL6, N=17). P values in panel c: one-way ANOVA with Dunett’s multiple comparisons test (post-hoc); panel d: P values according to Student’s t-distribution for Nc cor and hypergeometric testing for Num. panel f: one-way ANOVA with Tukey’s multiple comparisons test (post-hoc); comparisons between groups labeled in red are depicted in the bar graph. Asterisks indicate significance between groups (* P<0.05 to **** P<0.0001); All error bars correspond to standard deviation (Mean ± SD).

**Supplementary Figure 4:**
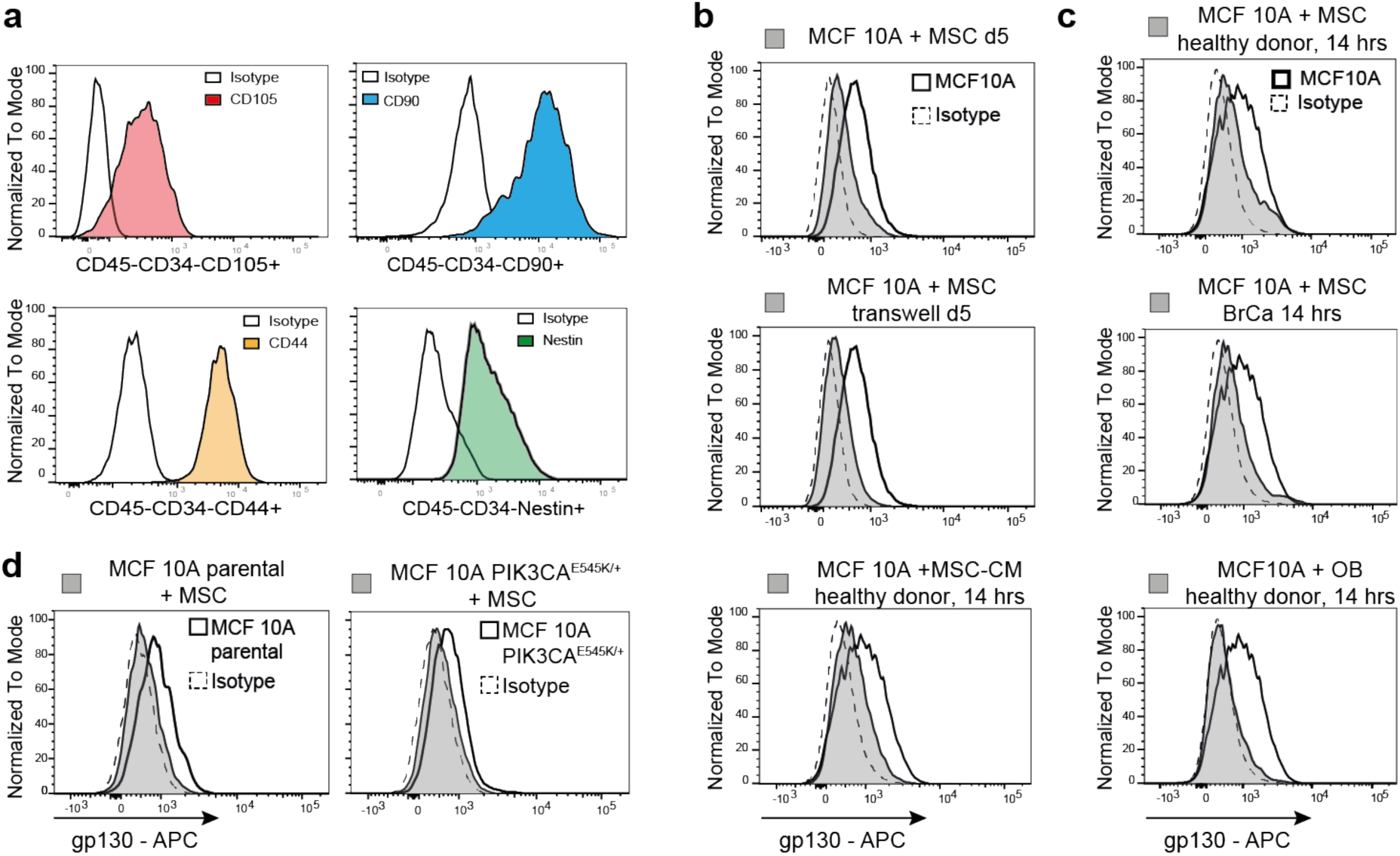
Surface expression of gp130 is down-regulated by soluble factors of bone marrow stromal cells. **a** MSCs isolated from bone marrow biopsies of healthy patients or patients with non-metastasized breast cancer were CD45^-^CD34^-^CD105^+^CD90^+^CD44^+^Nestin^+^. **b** gp130 surface expression of MCF 10A cells after five days of co-culture with MSCs or MSCs separated by a transwell or after 14 hrs of co-culture with MSC-conditioned medium (MSC-CM). **c** gp130 surface expression of MCF 10A cells after 14 hours of co-culture with MSCs or OBs from a healthy donor or breast cancer patient. **d** gp130 surface expression on isogenic MCF 10A cells without (MCF 10A parental) or with activating PIK3CA^E545K/+^ mutation cultured with MSCs for 5 days. **Panel b, c, d:** grey filled histograms indicate MCF 10A, MCF-7, or the isogenic cells MCF 10A parental and MCF 10A PIK3CA^E545K/+^ cells co-cultured with MSCs, OBs, MSC-CM or MSC separated by a transwell. Histograms with a thick black line indicate MCF 10A, MCF-7, or the isogenic cells MCF 10A parental and MCF 10A PIK3CA^E545K/+^ cells cultured alone and dashed histograms isotype control staining for gp130.

**Supplementary Figure 5:**
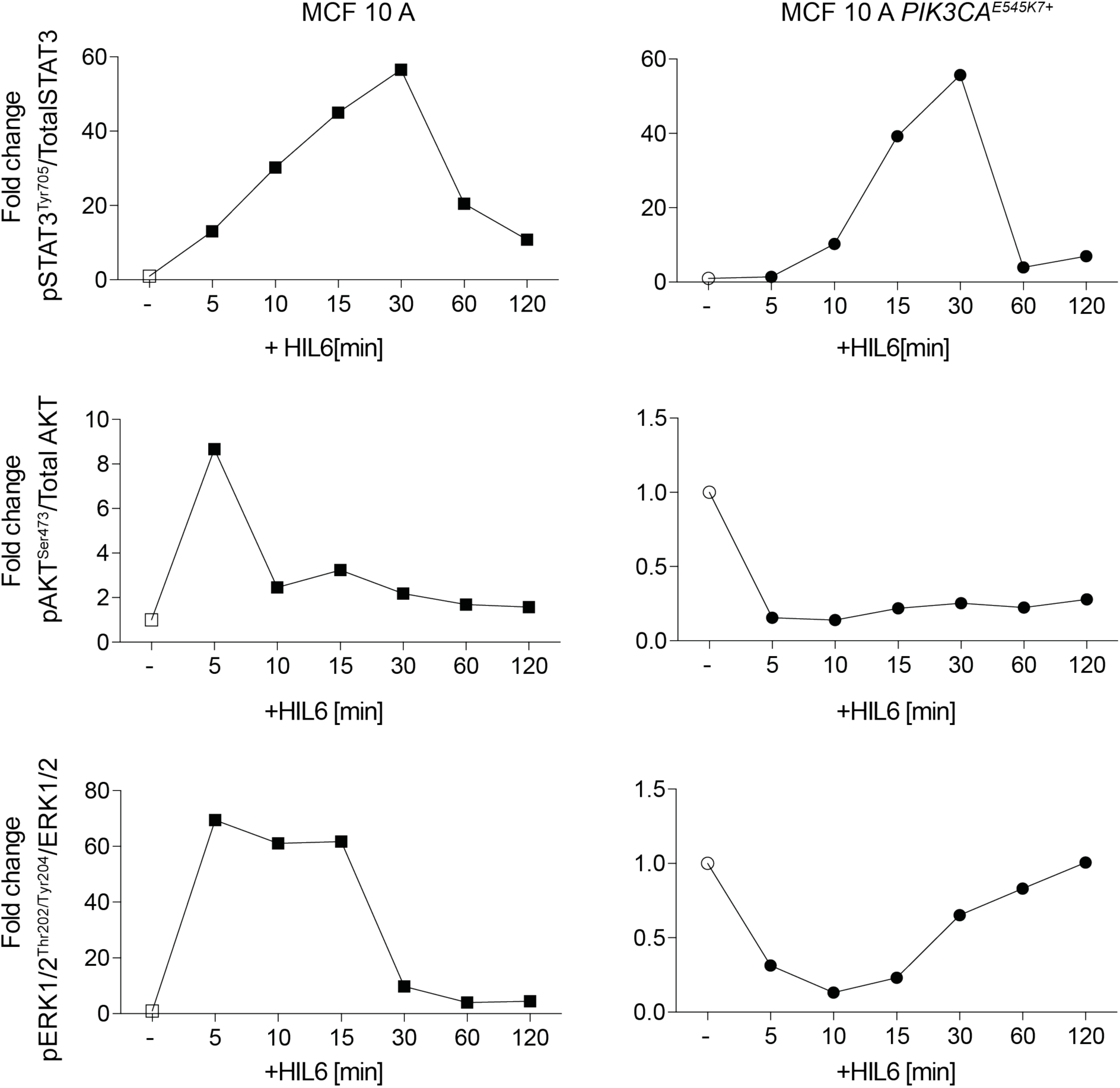
Activation of the JAK/STAT, PIK3CA/AKT and MAPK/ERK pathway by HIL6 stimulation. Quantification of western blot analyses (Fig. 7b) showing phosphorylation of STAT3^Tyr705^, AKT^Ser475^ and ERK1/2^Thr202/Tyr204^ in MCF 10A or MCF 10A *PIK3CA E545K/+* cells in the absence (open symbols) or presence of HIL6-stimulation (filled symbols). The signal from phosphorylated proteins and total proteins were normalized to *α*-tubulin before the ratio of phosphorylated proteins to total proteins was calculated. The graphs show the fold change in signal ratio over time relative to the respective control (unstimulated MCF 10A or MCF 10A *PIK3CA E545K/+*).

**Supplementary Figure 6:**
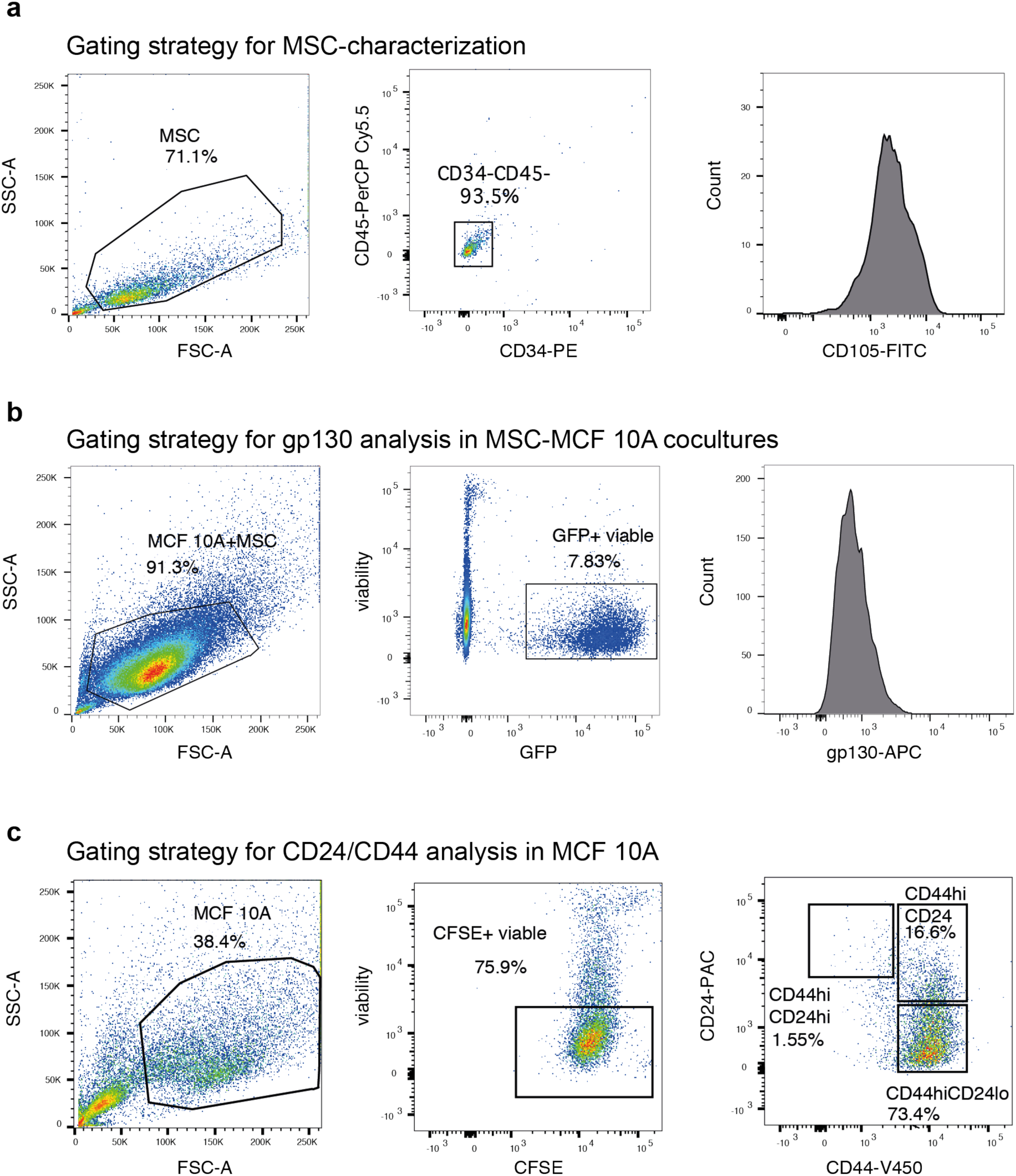
Gating strategies for flow cytometric analysis. **a** MSC-characterization; **b** gp130 analysis in MCF 10A-GFP cells co-cultured with MSCs, OBs or HUVECs; **c** CD24/CD44 analysis in CFSE-labeled MCF 10A cells.

## Notes

### Competing Interest Statement

CAK is in the scientific advisory board of HiberCell Inc, USA

